# Self-antigen disrupts cDC1 mediated antitumor responses

**DOI:** 10.64898/2026.06.25.734634

**Authors:** Victoria P Schuster, Kasidy Brown, Valentina Laverde, Naomi Berkowitz, Alyssa M Granados, Naoki Oshimori, John M Leech, Antonin Weckel, Tiffany C Scharschmidt, Megan K Ruhland

## Abstract

Conventional dendritic cells navigate complex tissues and sample peripheral antigens, balancing immune suppression and activation to achieve tissue homeostasis. However, it remains unclear how an individual dendritic cells reconciles co-incident signals from immunological opposing antigens within the tissue or tumor microenvironment, where tolerogenic self-antigen and immunogenic tumor antigen or microbes coexist. Here, using complementary *in vivo* fluorescent reporter systems, high-resolution imaging and endosomal profiling, we simultaneously tracked uptake, intracellular processing and cross-presentation of cutaneous self, tumor and microbial antigens in murine skin tissue, tumors and draining lymph nodes. We find that a substantial fraction of dendritic cells acquire antigen from multiple sources and that localization within endosomal compartments is dictated by antigen source. Notably, type 1 conventional dendritic cells that co-process self and tumor antigen represent a significant proportion of tumor antigen-bearing dendritic cells in the tumor and tumor draining lymph node. These dual-antigen loaded dendritic cells display a diminished capacity to prime tumor-specific CD8^+^ T cells and a marked reduction in tumor derived peptide presented on surface MHCI, while cross-priming of self-antigen specific T cells is significantly increased. These changes occur despite equivalent or greater tumor antigen uptake relative to self-antigen and high expression of surface MHCI and costimulatory molecules. Together, these data support a model in which multiantigen processing within dendritic cells can bias peptide loading away from tumor-derived epitopes, thereby limiting tumor-specific cross-priming. Modulating the antigenic context of the tumor microenvironment or endosomal routing after antigen uptake may therefore represent a strategy to restore effective dendritic cell-mediated antitumor immunity.

## INTRODUCTION

The proficiency of conventional dendritic cells (cDCs) to cross-present antigens from exogenous sources to CD8^+^ T cells enables dynamic regulation of immune activation and preservation of tolerance. Upon cDC sampling of peripheral tissues and associated antigenic pattern recognition receptor (PRR) signaling, antigens are recognized and processed for downstream cross-presentation through either the cytosolic or vacuolar pathway (1–3). In the cytosolic pathway, exogenous antigens undergo phagosomal escape, are degraded by the proteosome, and are then loaded onto major histocompatibility complex I (MHCI) in a transporter associated with antigen processing (TAP)-dependent manner (4–6). Within the vacuolar pathway, phagosomal maturation allows for antigenic peptide generation and MHCI loading in a TAP-independent manner (7–10). *In vitro* research supports a dependence on antigen-derived PRR signaling for appropriate phagosomal processing by antigen presenting cells (APCs) (11–15). Therefore, signaling induced by and inherent to the antigen provides a mechanism for tailoring immune responses to the peripheral tissue context (14,16–18). Within tissues, particularly barrier tissues, a milieu of host and foreign antigen sources can simultaneously provide context to cDCs within the same environment. Yet, our understanding of how concurrent PRR signaling impacts the vacuolar cross-presentation pathway to mediate immune activation or tolerance is limited to *in vitro* studies and lacks inclusion of cDCs with clinically relevant tissue sources for antigen context.

Within the cancer immunity cycle, cross-presentation of tumor-derived antigen facilitates the cytotoxic antitumor response (19,20). cDCs sample cell associated antigen at the tumor site, traffic to tumor draining lymph nodes (tdLNs), and present the processed antigen as peptides on MHCI. Prior work has established that the transcriptionally distinct type 1 conventional dendritic cell (cDC1) subset yields superior cross-presentation of tumor antigen and correlates with improved antitumor responses and survival in patients with solid tumors (21–24). This key role of cDC1s in the cancer immunity cycle has been elucidated from simplified model systems restricted to tracking tumor derived antigen (21,24,25). However, the complexity of the tumor microenvironment (TME) complicates our narrow understanding of cDC1-mediated tumor antigen cross-presentation. Spatial analyses have revealed heterogeneous activating and tolerizing antigen sources beyond tumor cells that comprise the TME, including non-cancerous, self-derived cells and microbes (26–28). Consequently, cDCs must regulate differential processing of non-tumor derived antigens from the TME to induce a successful tumor-specific CD8^+^ T cell response via cross-presentation. While some PRR signaling pathways within cDC1s have been identified to promote tumor cell associated cross-presentation (29–31), the direct implications of tumor cross-presentation by cDC1 sampling of additional TME antigen sources, such as non-cancerous self, alongside tumor antigen has yet to be determined.

In this study, we combine *in vivo* fluorescent model systems to track endosomal processing of self, tumor and microbial sourced antigens within cDCs. With these tools, we demonstrate that cDCs can process multiple TME antigens simultaneously yet autonomously through intracellular endosomal organization, and that a significant population of cDCs simultaneously process self and tumor antigen *in vivo*. In cDC1s, this leads to a suppression of tumor specific CD8^+^ T cell proliferation, revealing how cDC management of the heterogeneous TME antigens can alter antitumor immunity.

## RESULTS

### cDCs interact with multiple antigen sources in skin tissue, intracellularly sorting them into distinct compartments based on their source

Skin antigen-specific responses driven by cDC1 cross-presentation can lead to both tolerogenic and inflammatory immune responses (32–35). To begin investigating multiantigen handling within cDCs *in vivo*, we combined multiple fluorescent models to simultaneously track cDC uptake of previously defined tolerogenic skin self-derived material (33,34) and pathogenic skin microbe-derived material (35,36). We developed a K14-cre; lox-stop-lox-tdTomato (K14-tdTom) mouse line where trafficking of migratory cDCs that had taken up keratinocyte derived self-material (tdTom^+^ cDCs) could be identified within skin draining lymph nodes (dLNs) (Fig. S1A-S1B). Topical application of the common skin pathogen *Staphylococcus aureus* (*S. aur*) expressing ZsGreen (*S. aur*-ZsG) to the dorsal skin of (36) enabled us to generate an *in vivo* model system that allowed for fluorescent tracking of cDCs that had taken up material from either tolerogenic (skin, tdTom^+^) or inflammatory (*S. aur*, ZsG^+^) skin derived sources or both (tdTom^+^ZsG^+^) (Fig. 1A-B). Flow cytometry analysis of cDCs within the skin demonstrated that cDCs were able to take up either self-derived material (tdTom^+^), pathogen-derived material (ZsG^+^) or both self- and pathogen-derived material (tdTom^+^ZsG^+^) (Fig. 1B-C, Fig. S1C). The presence of a tdTom^+^ZsG^+^ cDC population highlighted the capability of skin cDCs to simultaneously handle material from different sources within the skin. To rule out artifacts driven by the fluorescent proteins, we topically applied the same microbe expressing mCherry (*S. aur*-mCh) to K14-cre; lox-stop-lox-ZsGreen mice (K14-ZsG) and found a similar uptake pattern (Fig. S1D). To visualize the intracellular antigen localization within individual cDCs, we sorted ZsG^+^mCh^+^ cDCs from skin of K14-ZsG mice topically associated with *S. aur*-mCh and performed live, confocal imaging (Fig. 1D). The antigen-bearing compartments were present as distinct puncta within the cDCs (Fig. 1E). On a per cell basis, pathogen^+^ puncta rarely colocalized with self^+^ puncta (Fig. 1E-F). Specifically, cDCs maintained separation of material derived from self-ZsG^+^ and *S. aur*-mCh^+^, resulting in few self^+^ *S. aur*^+^ puncta within the ZsG^+^mCh^+^These findings are c*in vitro* models (12,14).

**Figure 1.**
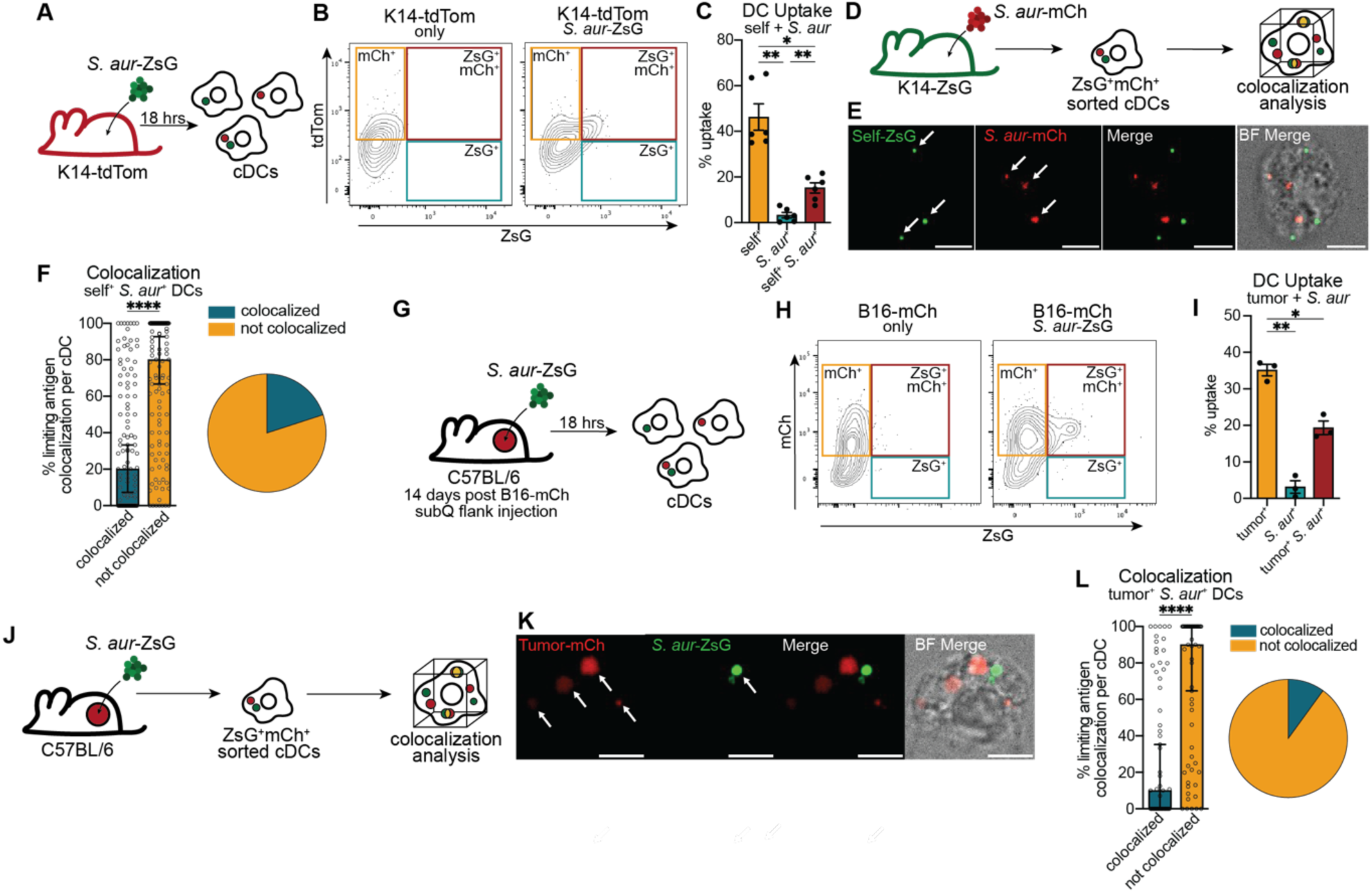
Dendritic cells interact with multiple antigens in tissues, sorting them into individual compartments based on antigen source. (A) Schematic of *S. aureus*-ZsGreen (*S. aur*-ZsG) application to K14-Cre; lsl-tdTomato (K14-Cre; lsl-tdTom) self reporter mice *in vivo*. (B) Representative flow cytometry plots of skin cDC uptake (live/CD45^+^/B220^-^, Ly6G^-^, NK1.1^-^, Ly6C^-^, CD90^-^/MHCII^+^, CD11c^+^) from uncolonized K14-Cre; lsl-tdTom skin only (left) or *S. aur*-ZsG colonized K14-Cre; lsl-tdTom (right). (C) Skin cDC uptake quantification of *S. aur*-ZsG colonized K14-Cre; lsl-tdTom. (D) Schematic of *S.aureus*-mCherry (*S. aur*-mCh) application to K14-Cre; lsl-ZsGreen (K14-Cre; lsl-ZsG) self reporter mice and endosomal analysis of skin cDCs by confocal imaging. (E) Representative images of self-ZsG and *S. aur*-mCh from sorted ZsG^+^mCh^+^ cDCs. (F) Median colocalized limiting antigen/cell vs not colocalized in ZsG^+^mCh^+^ (self^+^*S. aur*^+^) cDCs from (D, E). (G) Schematic of *S. aur*-ZsG colonization of wildtype mice injected with B16-mCherry (B16-mCh) tumors 14 days prior to colonization *in vivo*. (H) Representative flow cytometry plots of tumoral cDC uptake from uncolonized wildtype mice injected with B16-mCh tumors (top) or from wildtype mice with B16-mCh tumors colonized with *S. aur*-ZsG (bottom). (I) Tumoral cDC uptake quantification of *S. aur*-ZsG colonized wildtype mice injected with B16-mCh. (J) Schematic of *S. aur*-ZsG colonization of wildtype mice injected with B16-mCh tumors 14 days prior to colonization *in vivo* and endosomal analysis of tumoral cDCs by confocal imaging. (K) Representative images of *S. aur*-ZsG and B16-mCh from sorted ZsG^+^mCh^+^ cDCs. (L) Median colocalized limiting antigen/cell vs not colocalized in ZsG^+^mCh^+^ (tumor^+^*S. aur*^+^) cDCs from (J, K). (C) Experiments are from dorsal mouse skin of individual mice. Each point represents one mouse (n=6, 2 independent experiments). (D-L) Experiments are from pooled tissue from 6-12 mice. (I) Each point represents data from one pooled mouse experiment (n = 3, 3 independent experiments). (F, L) Data are shown as median ± SEM. Statistical analysis was performed using a two-tailed unpaired Mann-Whitney U test. Each point represents the percentage of colocalized antigen within a single cell from (F) 3 or (L) 2 independent experiments with cells sorted from the pooled tissue of 6-12 mice. (C, I) Data are shown as mean ± SEM. Statistical analysis was performed using RM one-way ANOVA, Geisser-Greenhouse correction, with Tukey’s multiple comparison test. *p < 0.05, **p < 0.01, ***p < 0.001, ****p < 0.0001.

### cDCs simultaneously process exogenous material from multiple sources within the TME

Considering the capacity of cDCs within skin tissue to manage multiple antigen sources concurrently, we next examined whether this phenotype was observed within the skin TME. Recent reports have shown that microbes serve as a clinically relevant, non-tumor antigen source within the skin TME (26,27,37). Therefore, we used the fluorescent B16F10 melanoma tumor model expressing mCherry (B16-mCh) alongside *S. aur*-ZsG to determine whether cDCs process multiple antigen sources simultaneously within the TME. 14 days post subcutaneous implantation of B16-mCh tumors into wildtype mice, *S. aur*-ZsG was applied to the overlaying unbroken skin (Fig. 1G). The following day, tumors were harvested, digested and stained for flow cytometry (Fig. 1H-I). While the majority of tumoral mCh^+^ cDCs had only taken up tumor-expressed mCh, the presence of ZsG^+^ cDCs and mCh^+^ZsG^+^ cDCs demonstrated the ability of cDCs to take up both tumor material and *S. aur*-derived material from the TME and adjacent skin tissue (Fig. 1H-I). We next sorted out the mCh^+^ZsG^+^ (tumor^+^*S. aur*^+^) cDCs from the TME and performed live cell imaging as before (Fig. 1J-K). Similar to the mCh^+^ZsG^+^ (self^+^*S. aur*^+^) cDCs, there was little colocalization observed between the tumor^+^ (mCh^+^) puncta and the *S. aur*^+^ (ZsG^+^) puncta within the tumor^+^*S. aur*^+^ cDCs (Fig. 1L). These results show that cDCs sample antigen sources beyond tumor antigen at the tumor site, a previously unidentified function of cDCs within the TME. The cDCs’ dynamic regulation of compartments bearing material from different antigen sources suggests an antigen source specific response, which could ultimately impact cDC maturation and antigen processing.

### Dendritic cell multiantigen processing correlates with increased maturation

The maturation state of cDCs directly determines the quality of downstream T cell priming, thereby biasing the response toward tolerance or productive activation. Under steady-state conditions, uptake of self-antigen does not result in cDC maturation, thus restricting autoimmune, self-specific T cell activation. How simultaneous uptake of multiple different antigen sources influences cDC maturation is unclear. To determine whether the uptake of multiple antigen sources impacts cDC maturation, we generated an *in vitro* fluorescent model for self-derived antigen by immortalizing a keratinocyte line (K14-ZsG-OVA cell line) derived from the neonatal skin of a K14-cre; lsl-ZsGreen; K14-OVA (K14-ZsG-OVA) mouse which has keratinocyte restricted expression of both ZsGreen and the model antigen ovalbumin (OVA). We confirmed ZsGreen expression by flow cytometry (Fig. S2A) (38). DC uptake of antigen from K14-ZsG-OVA cells was confirmed as was uptake of B16-ZsG-OVA (B16F10 expressing ZsGreen and OVA) tumor cells (Fig. S2A) (39). To recapitulate our *in vivo* systems that allow for multiantigen source uptake, we cocultured 1x10^5^ bone marrow derived dendritic cells (BMDCs) with 1x10^5^ mitomycin C induced apoptotic K14-ZsG-OVA or B16-ZsG-OVA cells (1:1) and 1x10^6^ CFUs of *S. aur*-mCh (MOI of 10) for 4 hours. *S. aur*-mCh microbes were heat-killed prior to coculture to prevent BMDC infection and culture overgrowth. To determine the impact on cDC maturation following multiantigen source uptake, cocultures were collected and BMDCs were stained for flow cytometry using a cDC phenotyping antibody panel (Fig. S2B). Compared to BMDCs that had taken up *S. aur* material only, co-uptake of either tumor and *S. aur* (Fig. S2C) or self and *S. aur* (Fig. S2D) derived material from the same coculture environment led to significantly increased expression of costimulatory molecules CD80 and CD86 (Fig. S2C-D). Co-uptake of *S. aur* and either tumor or self-derived antigens also led to increased IL-12 and IL-10 cytokine production compared to BMDCs that had only taken up *S. aur* (Fig. S2E-F). Production of both IL-10 and IL-12 was significantly greater in the self^+^*S.aur*^+^ population compared to the only self^+^ population (Fig. S2F) whereas only IL-12 showed a significant difference when directly comparing tumor^+^ versus tumor^+^*S.aur*^+^ BMDCs. These data suggest that within a given condition, both costimulation and cytokine production from DCs are dependent on the identity of the antigen source, with uptake of cell-derived antigen (tumor or self) leading to increased levels of both IL-10 and IL-12 compared to *S. aur*^+^ alone. Surface levels of MHCI trended towards highest increased expression in BMDCs that had taken up either tumor and *S. aur* (Fig. S2G) or self and *S. aur* (Fig. S2H) though values did not reach statistical significance in the tumor cocultures. Taken together, these data indicate that the uptake of antigens from multiple sources generally leads to increased cDC maturation. These findings parallel research describing increased cDC maturation both in the TME and tdLN when strong PRR signaling is induced in tumor antigen-bearing cDCs (18).

### DC endosomal processing of antigen is dependent on antigen source

Given our finding that skin- and TME-derived cDCs organize antigens intracellularly based on antigen source, we next sought to determine whether antigen source identity impacts endosomal processing in environments with multiantigen uptake. To determine whether endosomal cargo impacts downstream processing, we modified published *in vitro* phagoFACS techniques ⍰(14,29,40)⍰. Coculture assays were set up as in Fig. S2B, followed by CD11c-positive enrichment and lysis. Gradient isolation was used to purify endosomes from the lysed *in vitro* coculture assays (Fig. 2A). Purified endosomes were surface stained with endosomal pathway markers to define endosomal processing dynamics. Size calibration beads facilitated the identification of endosomes via flow cytometry (Fig. 2B). We found no difference in endosomal association of the lysosomal marker LAMP1 based on the identity of the cargo, inferring little impact of endosomal antigen identity on ⍰(11,41)⍰ (Fig. S2I). Comparatively, the marker for the late endosomal compartment, Rab7, was only found on endosomes harboring *S. aur*, but not tumor nor self-material loaded endosomes, independent of the coculture conditions (Fig. S2J). These data suggest that while *S. aur*-bearing endosomes may mature into late endosomes (e.g. Rab7^+^ endosomes) at a greater frequency than cell-derived material overall, antigen source does not appear to dictate lysosomal fate (e.g. LAMP1^+^). To determine the impact of antigen source on endosomal processing for cross-presentation, we stained for EEA1 and Rab5 which have been previously used to identify “sorting endosomes” associated with cross-presentation ⍰(42,43)⍰. We found that a significantly lower frequency of endosomes harboring either self- or tumor-derived material stained for cross-presentation associated markers compared to endosomes bearing *S. aur* derived material (Fig. 2C-2E). In agreement with the EEA1 and Rab5 association, only *S. aur* endosomes were found to be enriched in intraendosomal MHCI while endosomes harboring either tumor or self from the same coculture conditions did not efficiently recruit MHCI to the antigen-bearing endosome (Fig. 2F-2H). To determine if the endosomal processing observed was inherent to the antigen source or dependent on the coculture conditions, we independently cultured BMDCs with the individual antigen sources (*S. aur*, tumor or self). Similar to our previous observations, we found little association between lysosomal LAMP1-positivity and endosomal contents (Fig. S2K), but observed coordinate enrichment of the late endosomal marker Rab7 (Fig. S2L) and cross-presentation associated EEA1, Rab5 and MHCI (Fig. 2I-2K) on endosomes harboring *S. aur* derived material. Together, these data indicate that the antigen source can impact endosomal processing, particularly processing for cross-presentation. Further, these data suggest that cDC intracellular structural organization of antigen and differential endosomal processing may serve as a mechanism for maintaining proper immune reactivity within tissues where both tolerizing and activating antigens are present.

**Figure 2.**
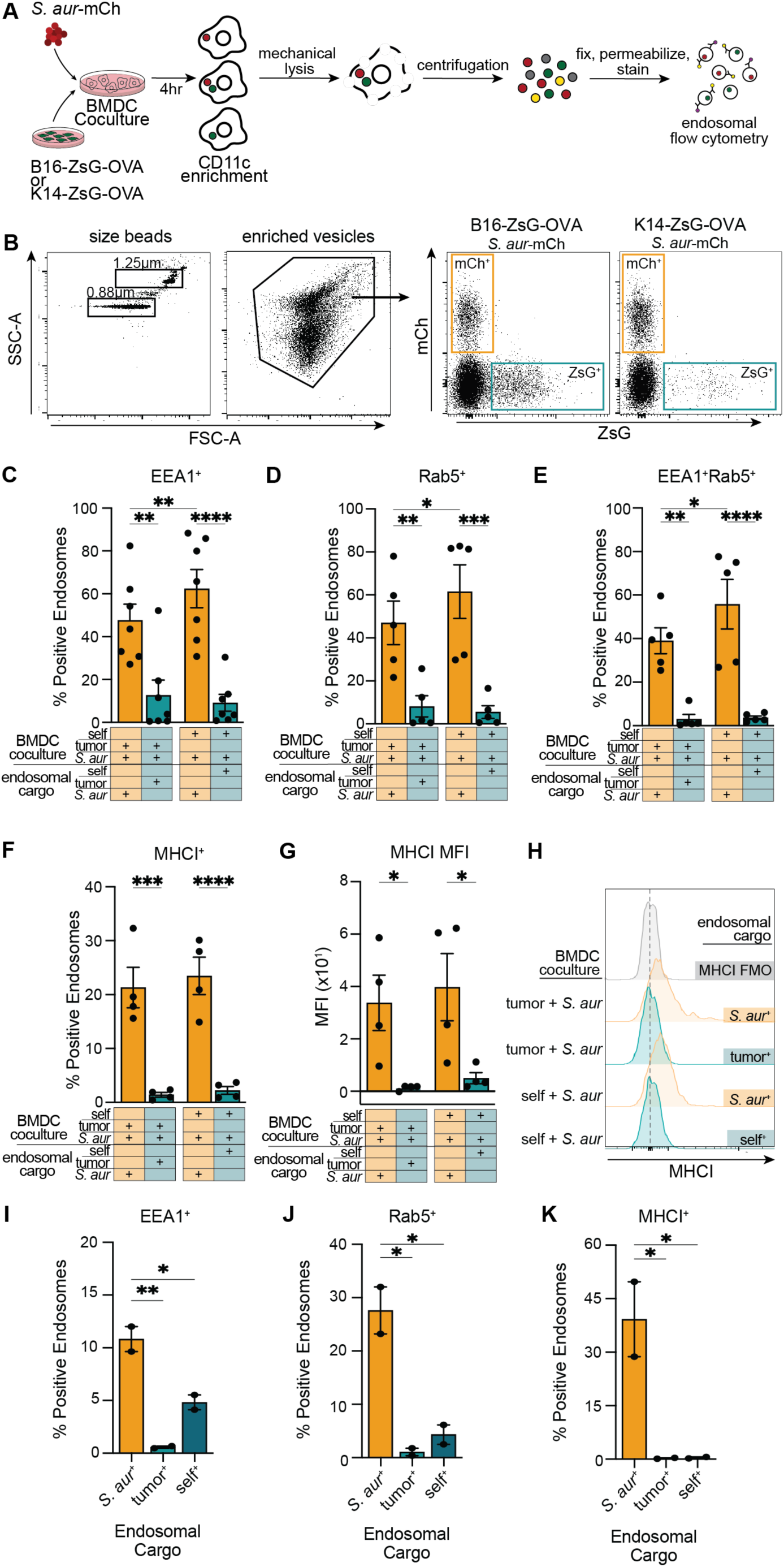
DC phenotype and endosomal processing is dependent on antigen source. (A) Schematic for endosomal phenotyping from BMDC coculture with tumor (B16-ZsG-OVA) or self (K14-ZsG-OVA) and *S. aur*-mCh. (B) Representative flow cytometry plots of isolated endosomes from (left) single *S. aur*-mCh (middle) dual B16-ZsG-OVA and *S. aur*-mCh or (right) K14-ZsG-OVA and *S. aur*-mCh antigen pulsed BMDCs. (C-F) Percent (C) EEA1^+^ (D) Rab5^+^ (E) EEA1^+^, Rab5^+^ (F) MHCI^+^ endosomes based on endosomal cargo from BMDCs pulsed with tumor and *S. aur* or self and *S. aur*. (G) MFI and (H) histograms of MHCI of endosomes based on endosomal cargo from BMDCs pulsed with tumor and *S. aur* or self and *S. aur*. (I-K) Percent (I) EEA1^+^ (J) Rab5^+^ (K) MHCI^+^ of endosomes from BMDCs pulsed with single antigen sources. (C-G) Data are shown as mean ± SEM. Statistical analysis was performed using RM two-way ANOVA with matched values. Post hoc comparisons were conducted using Fisher’s LSD test. Each point represents data from an individual BMDC experiment (n = 4-7, 4-7 independent experiments). (I-K) Data are shown as mean ± SEM. Statistical analysis was performed using one-way ANOVA with Tukey’s multiple comparison test. Each point represents data from an individual BMDC experiment (n = 2, 2 independent experiments). *p < 0.05, **p < 0.01, ***p < 0.001, ****p < 0.0001.

### DCs take up self and tumor antigen, sorting them into the same endosomal compartment

Consistent patterns in endosomal sorting, maturation, and processing of self and tumor antigens in relation to *S. aur* suggested that self and tumor sources may be handled similarly within complex tissue environments. To test this, we implanted B16-mChOVA tumor cells subcutaneously into the flanks of K14-ZsG mouse to enable concurrent tracking of tumor (mCh^+^) and self (ZsG^+^) derived material within cDCs. Both tumors and tdLNs were harvested, digested and processed for flow cytometry analysis of cDCs (Fig. 3A). Surprisingly, we found that not only are cDCs in both the TME and tdLNs capable of harboring both tumor and self-derived antigen concurrently (Fig. 3B-3D), but that the ZsG^+^mCh^+^ population represented a significant proportion of the antigen-bearing cDC population at these critical tissue sites. Even in our model system where self-antigen is singularly defined by keratinocyte origin, the proportion of self^+^tumor^+^ cDCs in the TME still represented the largest ratio of cDCs that have taken up fluorescently labeled antigens (Fig. 3C, 3E). In the tdLNs, we also found a large population of antigen^+^ migratory cDCs harboring both tumor and self (Fig. 3D, 3E). Self^+^ cDCs represented the largest proportion of antigen^+^ migratory cDCs in the tdLN (Fig. 3C-E). This is consistent with the constitutive nature of K14-Cre and the expression of ZsGreen throughout the entire (tumor uninvolved) skin of K14-ZsG animals which can drain to the same LN as the tumor. Prior research has also shown that skin antigen loaded migratory cDCs traffic to LNs both at steady state and under inflammatory conditions (39,44). Regardless, self^+^tumor^+^ cDCs represent about half, if not more, of the total proportion of cDCs in the tdLN that have taken up tumor antigen (Fig. 3D, 3F).

**Figure 3.**
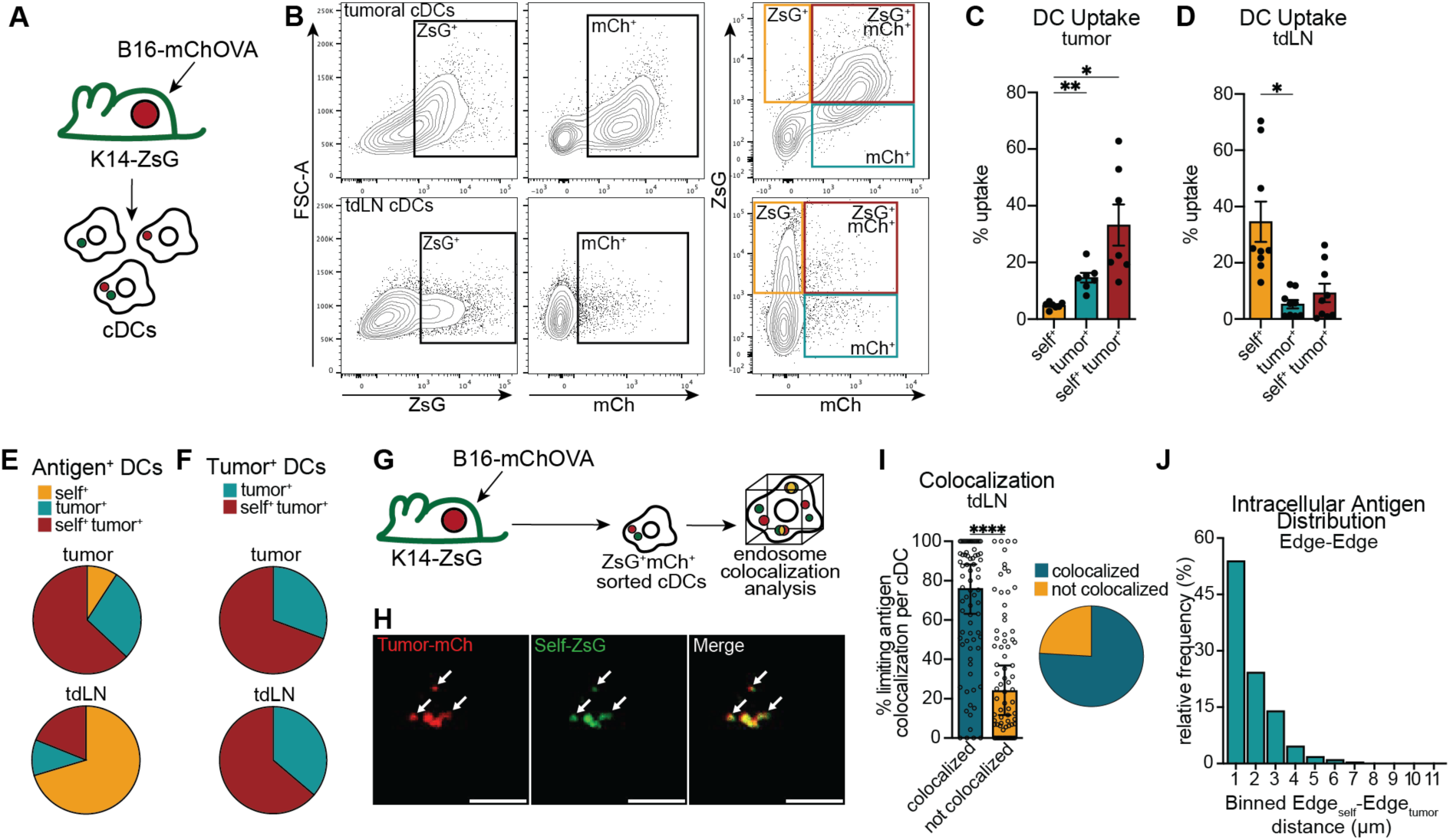
DCs take up self antigen alongside tumor antigen, colocalizing them into the same endosomal compartment for processing. (A) Schematic of mouse model to track tumor and self antigen uptake by cDCs *in vivo*. (B) Flow plots for uptake of cDCs within the (top) TME and (bottom) tdLNs gated on cDCs. (C-D) Quantification of percent uptake in (C) tumor tissue and (D) tdLNs. (E-F) Ratio of cDC uptake by cDCs that have taken up (E) self or tumor antigen or (F) tumor antigen in (top) TME) and (bottom) tdLNs from (C-D). (G) Schematic for colocalization analysis of tumor and self antigen by cDCs *in vivo*. (H) Representative images of tumor^+^self^+^ cDCs from tdLN. (I) Quantification of limiting antigen colocalization of tumor and self antigen within the tdLN cDCs. (J) Binned edge-to-edge distance analysis between intracellular antigens of each antigen source in tdLN. Data are shown as mean distance between EdgeA-EdgeB of antigens within an individual cell from (I). (C-D) Data are shown as mean ± SEM. Statistical analysis was performed using RM one-way ANOVA, Geisser-Greenhouse correction, with Tukey’s multiple comparison test. Each point represents data from an individual mouse (n = 6, 2 independent experiments). (I) Data are shown as median ± SEM. Statistical analysis was performed using a two-tailed unpaired Mann-Whitney U test. Each point represents the percentage of colocalized antigen within a single cell, 3 independent experiments with cells sorted from the pooled tissue of 8-12 mice. *p < 0.05, **p < 0.01, ***p < 0.001, ****p < 0.0001.

To evaluate endosomal compartmentalization of self and tumor antigen within individual cDCs, we used fluorescent microscopy to quantify colocalization of antigens in sorted self^+^tumor^+^ cDCs in the tdLNs (Fig. 3G). 3D imaging analysis revealed a high rate of colocalization of these two antigen sources into the same endosomal compartments within the self^+^tumor^+^ cDCs (Fig. 3H-I). Spatial distance analysis of the antigens confirmed close intracellular proximity and endosomal organization within the individual cDCs (Fig. 3J, S3A). To determine the impact of OVA in our system, we repeated these experiments using B16-mChOVA tumors in the K14-ZsG-OVA mouse strain and found no appreciable difference in colocalization of self and tumor-expressed antigen (Fig. S3C-D) suggesting that a pre-established central tolerance to OVA does not impact endosomal sorting of tumor sourced antigen. Critically, these data suggest that self^+^tumor^+^ cDCs represent a substantial proportion of the cDC population.

### Concurrent uptake of self-antigen by cDCs diminishes cross-priming of tumor antigen

The crucial role of tumor antigen bearing cDCs in cross-priming anti-tumor CD8^+^ T cells is well established (24,39,45). However, how the presence of self-antigen within tumor antigen-loaded cDCs impacts the ability to prime anti-tumor T cells is currently unknown. Co-processing of self and tumor antigens within individual cDCs could influence which peptide source is routed for MHCI loading and presentation, with potential consequences for self and/or tumor antigen selection. To test this possibility directly, we examined whether concurrent uptake of tolerogenic self-antigen alters cDC cross-priming of tumor-specific CD8^+^ T cells.

We leveraged our fluorescent protein systems with the model antigen OVA together with CD8^+^ T cells expressing an OVA-specific T cell receptor (TCR). OT-I transgenic animals provided a source of CD8^+^ T cells specific for the OVA-derived SIINFEKL peptide presented on MHCI. This system allowed us to specifically quantify the efficiency of anti-tumor cross-priming upon coculture with cDCs sorted by uptake from B16-mChOVA tumors implanted in K14-ZsG mice (Fig. 4A). OT-I T cells labeled with violet proliferation dye were cocultured with cDCs sorted by antigen uptake at a 1:10 cDC:OT-I ratio for 72 hours. T cells were then harvested for antibody staining to determine proliferation and phenotyping via flow cytometry. While cDCs sorted from the TME that had taken up only tumor-expressed OVA antigen were capable of inducing proliferation of OT-I T cells, this response was significantly diminished when T cells were stimulated by cDCs that had also taken up self-antigen (Fig. 4B-C, S4A). We evaluated the T cells for functional markers and found that, despite inducing less proliferation, self^+^tumor^+^ cDCs elicit significantly greater PD-1 expression in activated T cells compared to tumor^+^ cDCs (Fig.4E). However, the CD8^+^ T cell response showed no dysfunction in effector cytokine production upon stimulation by self^+^tumor^+^ cDCs, compared to tumor^+^ cDCs (Fig. 4F). These data suggested that uptake of self material negatively impacts the ability of cDCs to induce proliferation of tumor-specific CD8^+^ T cells.

**Figure 4.**
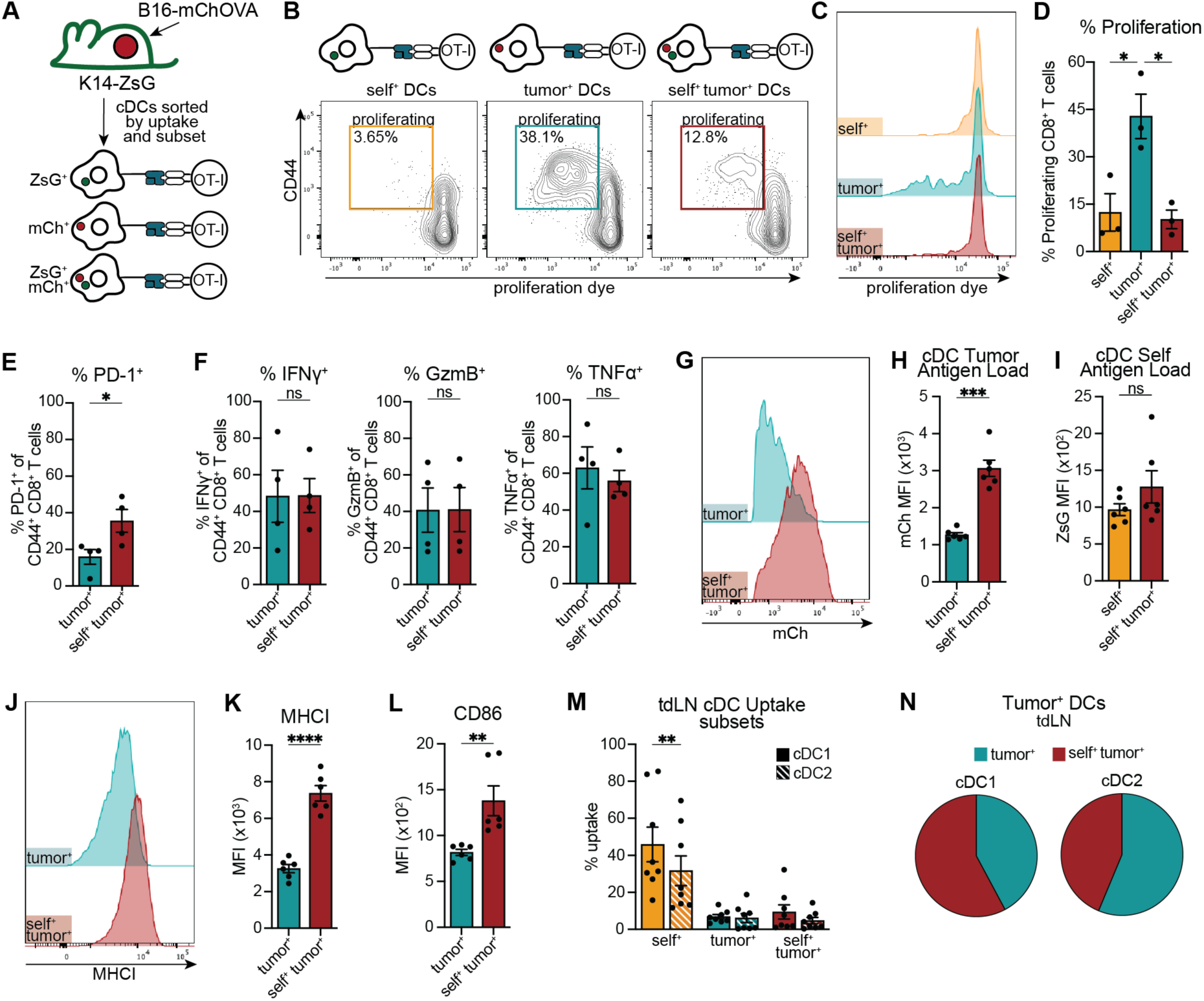
DC simultaneous processing of tumor and self impacts priming of tumor-specific CD8 T cells. (A) Schematic of tumor-specific OT-I CD8^+^ T cell stimulation assay with cDCs based on their antigen uptake. (B-C) Representative (B) proliferation graphs, (C) proliferative histograms of OT-Is from cDCs that have taken up either self, tumor, or both tumor and self. (D-E) Quantification of (D) percent proliferative and (E) percent PD-1^+^ OT-Is cocultured with DCs sorted based on uptake. (F) Percent IFN-γ^+^, GzmB^+^, and TNFa^+^ of CD44^+^ CD8 T cells following coculture with sorted DCs. (G) Representative histogram and (H-I) quantification of uptake by MFI of (G-H) tumor antigen or (I) self antigen by intratumoral DCs. (J) Representative histogram and (K) MFI of MHCI from tumor^+^, or tumor^+^self^+^ intratumoral DCs. (L) MFI of CD86 from tumor^+^, or tumor^+^self^+^ intratumoral DCs. (M) Quantified uptake and (N) ratio of uptake by antigen^+^ migratory cDC1s (mDC1s) and migratory cDC2s (mDC2s) from the tdLNs. (D-F) Data are shown as mean ± SEM. Statistical analysis was performed using (D) one-way ANOVA, with Tukey’s multiple comparison test or (E-F) two-tailed unpaired t-test. Each point represents the mean data from an individual experiment set up with 8-12 mice pooled per experiment. (D) n = 3, 3 independent experiments or (E-F) n = 4, 4 independent experiments. (H-I, K-L) Data are shown as mean ± SEM. Statistical analysis was performed using two-tailed paired t-test. Each point represents an individual mouse. n = 6, 2 independent experiments. (M-N) Data are shown as mean ± SEM. n = 8, 2 independent experiments. (M) Statistical analysis was performed using two-way RM ANOVA with matched values (stacked and across rows) and Geisser-Greenhouse correction. Post hoc comparisons were conducted using Šídák’s multiple comparisons test. Each point represents an individual mouse. *p < 0.05, **p < 0.01, ***p < 0.001, ****p < 0.0001.

It is possible that uptake of self material limits the amount of tumor material that a cDC can concurrently uptake. This could result in less tumor antigen availability within the cell and consequently reduce anti-tumor cross-priming. Surprisingly, we found that the reduction in T cell proliferation was not associated with a reduced amount of per cell tumor material (as determined by mean fluorescence intensity, MFI). In fact, self^+^tumor^+^ cDCs had taken up significantly more tumor-derived material than their tumor^+^ cDC counterparts isolated from the same TME (Fig. 4G-H). We observed no significant difference in the uptake of self-derived material when comparing self^+^ versus self^+^tumor^+^ cDCs within the TME (Fig 4I). Consequently, the ratio of tumor to self-derived material within the self^+^tumor^+^ cDCs was significantly higher than the ratio between the tumor^+^ cDCs and self^+^ cDCs (Fig. S4B). Uptake patterns within migratory cDCs of tdLNs similarly showed neither a significant reduction of tumor material in self^+^tumor^+^ cDCs compared to tumor^+^ cDCs nor changes in self material observed (Fig. S4C-S4D).

Given the significant differences in T cell stimulation based on the type of antigen load, we next asked if the presence of self-antigen within tumor-antigen bearing cDCs impacted cell maturation as measured by surface expression of costimulatory molecules and MHCI, both of which are necessary for optimal cross-priming (46–50). cDC phenotyping by flow cytometry revealed a significant increase in both surface MHCI and costimulatory molecule expression in self^+^tumor^+^ cDCs compared to tumor^+^ cDCs (Fig. 4J-4L, S4E-S4F). These findings were consistent with our earlier *in vitro* maturation analysis (Fig. S2B-S2H) and suggest that reduced maturation of self^+^tumor^+^ cDCs is not the driver of impaired T cell activation by this cDC population.

Given that neither the amount of tumor material within the cDCs nor their maturation explained the reduced T cell priming capacity of self^+^tumor^+^ cDCs, we next considered whether this difference stemmed from the subset of cDCs that had taken up tumor or tumor and self-antigen within the TME. For this we used flow cytometry to examine the cDC subsets within the TME and tdLN based on antigen-load. Within the tumor, we found that both cDC1 and type 2 conventional DC (cDC2) subsets acquired self and tumor derived material to varying degrees. A larger proportion of the cDC1s had taken up tumor alone whereas self^+^tumor^+^ cells represented the largest proportion of the cDC2 subset (Fig. S4G-J). However, when we profiled cDCs within the tdLN, which is the anatomical site of naïve CD8^+^ T cell priming, we found that self^+^tumor^+^ cDC1s and self^+^tumor^+^ cDC2s were a large proportion of tumor antigen-loaded cDCs. In fact, more than half of all tumor antigen bearing cDC1s and nearly half of cDC2s in the tdLN also contained self material (Fig. 4M-N). Given their abundance and impaired ability to stimulate T cells, these data indicate that self^+^tumor^+^ cDC1s and/or cDC2s may have consequential impacts on downstream T cell priming and warrant further consideration in the context of anti-tumor immunity.

### Simultaneous processing of self and tumor by cDC1s suppresses tumor-specific CD8^+^ T cell proliferation

In order to better understand relative contributions of cDC1s versus cDC2s to antigen-load dependent differences in CD8^+^ T cell priming, we performed T cell stimulation assays using either cDC1s or cDC2s with qualitatively different antigen loads. We cocultured OT-I CD8^+^ T cells with either migratory cDC1s or cDC2s sorted by uptake (tumor^+^ or self^+^tumor^+^) from the tdLNs of K14-ZsG mice implanted with B16-mChOVA (Fig. 5A, S5A). Self^+^tumor^+^ cDC1s failed to yield optimal tumor-specific CD8^+^ T cell proliferation compared to their tumor^+^ cDC1 counterparts (Fig. 5B-D), mirroring the phenotype we observed in bulk (cDC1 and cDC2 combined) TME-derived cDC cocultures (Fig. 4A-E). Intracellular cytokine staining again indicated that the function was similar between proliferating T cells stimulated with either tumor^+^ or self^+^tumor^+^ mDC1s (Fig. 5E, S5B). In contrast to self^+^tumor^+^ cDC1s, the ability of migratory cDC2s to drive tumor-specific CD8^+^ T cell activation was independent of the type of antigen taken up, with self^+^tumor^+^ cDC2s leading to similar proliferation compared to their tumor^+^ cDC2 counterparts (Fig. 5F-H). The profile of the activated T cells following coculture was also unchanged regardless of cDC2 uptake (Fig. S5C). Thus, the suppression of tumor-specific CD8^+^ T cell cross-priming upon self-antigen uptake is limited to cDC1s (Fig. 5I). Consistent with our earlier findings, acquisition of self-antigen was not associated with a decrease in tumor antigen uptake (Fig. 5J), surface MHCI expression (Fig. 5K) or costimulatory molecule expression (Fig. S5D) within the cDC1s or cDC2s within tdLNs, resembling their respective subsets within the TME (Fig. S5E-F). Phenotyping of the cDC2 subset was most similar to the bulk cDC phenotyping (Fig. 4J-L, S4E-F) as expected given the known relative proportions of cDC1:cDC2 (Fig. S5A). Take together, these data suggested that although cDC1s are known to drive superior CD8^+^ T cell responses to tumor antigen compared to cDC2s (21,24,25), uptake of self-derived material leads to a cDC1-specific reduction of T cell cross-priming. This exposes a critical gap in the understanding of how the uptake of self-derived material by cDC1s impacts the antitumor response.

**Figure 5.**
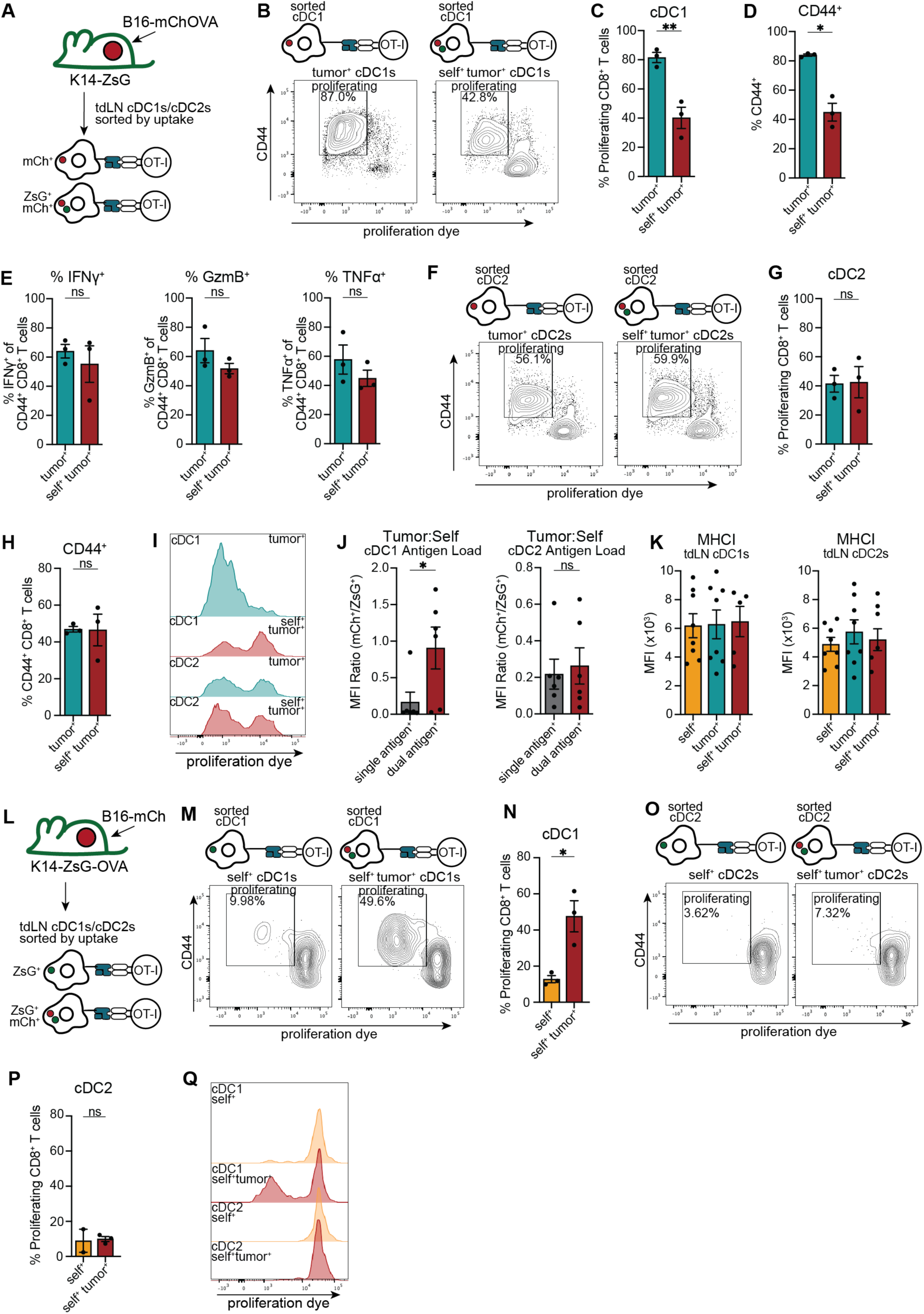
cDC1 processing of self and tumor depresses antitumor CD8 T cell proliferation. (A) Schematic of tumor-specific OT-I CD8^+^ T cell stimulation assay with migratory cDC1s (mDC1s) or cDC2s (mDC2s) from tdLNs based on their antigen uptake. (B,F) Representative proliferation graph and (C,G) quantification of percent proliferative OT-Is cocultured with (B-C) mDC1s or (F-G) mDC2s that are either tumor^+^ or tumor^+^self^+^. (D, H) Percent CD44^+^ OT-Is following coculture with (D) mDC1s or (H) mDC2s that are tumor^+^ or tumor^+^self^+^. (E) Percent IFNg^+^ (left), GzmB^+^ (middle), and TNFa^+^ (right) of CD44^+^ CD8 T cells following coculture with sorted mDC1s. (I) Histogram of proliferation by DC subsets that are either tumor^+^ or tumor^+^self^+^. (J) MFI ratio of tumor antigen to self antigen uptake by tdLN mDC1 (left) and mDC2 (right) subsets that have either taken up either tumor or self or both tumor and self antigen. (K) MFI of MHCI of mDC1s (left) and mDC2s (right) based on antigen uptake in tdLN. (L) Schematic of self-specific OT-I CD8^+^ T cell stimulation assay with cDC subsets from tdLNs based on their antigen uptake. (M,O) Representative proliferation graph and (N,P) quantification of percent proliferative OT-Is cocultured with (M-N) mDC1s or (O-P) mDC2s that are either self^+^ or tumor^+^self^+^. (Q) Histogram of proliferation by DC subsets that are either self^+^ or tumor^+^self^+^. (C-E, G-H, N, P) Data are shown as mean ± SEM. Statistical analysis was performed using (C, G, N, P) Statistical analysis was performed using a one-way ANOVA. Post hoc comparisons were conducted using Dunnett’s test, comparing each column mean to tumor^+^self^+^ mean. (D-E, H) two-tailed paired t-test. Each point represents the mean data from an individual experiment set up with 14-18 mice pooled per experiment. (D) n = 2-3, 3 independent experiments. (J, K) Data are shown as mean ± SEM. n = 8, 2 independent experiments. Each point represents an individual mouse. Statistical analysis was performed using (J) two-tailed paired t-test or (K) RM one-way ANOVA, Geisser-Greenhouse correction, with Tukey’s multiple comparison test. *p < 0.05, **p < 0.01, ***p < 0.001, ****p < 0.0001.

Given the presence of both self and tumor material in dual-loaded cDC1s, we hypothesized that reduced tumor antigen presentation may be the result of increased self-antigen presentation. To test this, we assessed whether dual-uptake by cDC1s resulted in differences in self-antigen cross-presentation. We transferred the source of model antigen (OVA) within the system from tumor to self. A strength of this approach is that by moving the OVA source rather than changing the model antigen, we can evaluate cross-priming efficiency, irrespective of differences in peptide:MHCI or peptide:MHCI:TCR binding affinities inherent to a given model antigen. We quantified the self-specific response by coculturing OT-Is with cDC1s or cDC2s from tdLNs of K14-ZsG-OVA mice implanted with B16-mCh tumors sorted by self^+^ or self^+^tumor^+^ uptake (Fig. 5L). Surprisingly, self^+^tumor^+^ cDC1s elicited increased self-specific CD8^+^ T cell proliferation, compared to self^+^ cDC1s from the same environment (Fig. 5M-5N, 5Q). The CD8^+^ T cells cocultured with self^+^tumor^+^ mDC2s displayed no change in proliferation compared to self^+^ mDC2s (Fig. 5O-5Q), recapitulating the response when the model antigen is sourced from the tumor (Fig. 5F-5G). Once more, the functional profile of these activated CD8^+^ T cells remained unchanged by the antigen uptake profile of either cDC subset (Fig. S5G-H). Taken together, these data indicate that co-uptake of self and tumor by cDC1s, but not cDC2s, amplifies self-specific CD8^+^ T cell priming, while concurrently blunting cross-priming of the antitumor T cell response.

### Self uptake disrupts antitumor T cell response by decreasing tumor derived peptide:MHCI and shifting endosomal processing

Prior *in vitro* work showed that antigen cross-presentation is dictated by the antigen source, though the impact of other antigens within the DC on the outcome of this process was not explored (12). Our findings suggest that there is a differential impact of dual-antigen uptake on the priming of self- versus tumor-specific T cells regardless of endosomal analyses showing that self and tumor material are preferentially co-housed within the same endosomes. These data suggest that cross-presentation of both tumor and self-antigen by cDCs is impacted by simultaneous processing of these antigens *in vivo* and raise the question of whether antigen selection for MHCI loading is biased for one antigen source over another. To test whether this simultaneous processing is ultimately impacting which antigen is selected for surface presentation by the self^+^tumor^+^ cDCs, we developed a sensitive bioluminescence assay that would enable us to quantify surface SIINFEKL:MHCI (SIIN:MHCI) complexes from *in vivo* antigen-loaded cDCs. Specifically, we used B3Z T cell hybridoma cells that contain a *LacZ* gene driven by the IL2 promoter and a TCR that specifically recognizes the SIIN:MHCI complex (51). Of note, B3Z stimulation does not require costimulation and thus provides a direct readout of TCR:pMHC engagement. Given that anti-H2K^b^-SIIN antibodies for flow cytometry staining of primary DCs can be artifact-prone, this assay enabled an antibody-independent means for measuring SIIN:MHCI complexes on cDCs. We first evaluated the specificity and sensitivity of B3Z stimulation by culturing the cells with increasing concentrations of either SIIN:MHCI or SIY:MHCI monomers (Fig. S6A). The validated SIIN:MHCI specificity of the monomer titrations provided a reference standard for quantification of presented peptide on cDCs within each B3Z stimulation experiment. To determine if the amount of tumor-derived peptide:MHCI on the cell surface was being impacted in self^+^tumor^+^ cDCs, we sorted tumor^+^ and self^+^tumor^+^ cDC subsets from the tdLNs of K14-ZsG mice injected with B16-mChOVA and performed a B3Z stimulation assay (Fig. 6A). We found that cDC1 uptake of self in addition to tumor antigen led to significantly less stimulation of B3Z T cells compared to cDC1 uptake of tumor alone (Fig. 6B). This was only seen with cDC1s, as the stimulation of B3Z T cells by tumor^+^ and self^+^tumor^+^ cDC2s remained similar (Fig. 6B). Using the monomer standard curve, we found that the number of tumor-derived peptide:MHCI on self^+^tumor^+^ cDC1s was less than half of that observed on tumor^+^ cDC1s (Fig. 6C-D). These data indicate that the decrease in cross-priming of tumor-specific CD8^+^ T cells by self^+^tumor^+^ cDC1s is driven by reduced tumor antigen cross-presentation.

**Figure 6.**
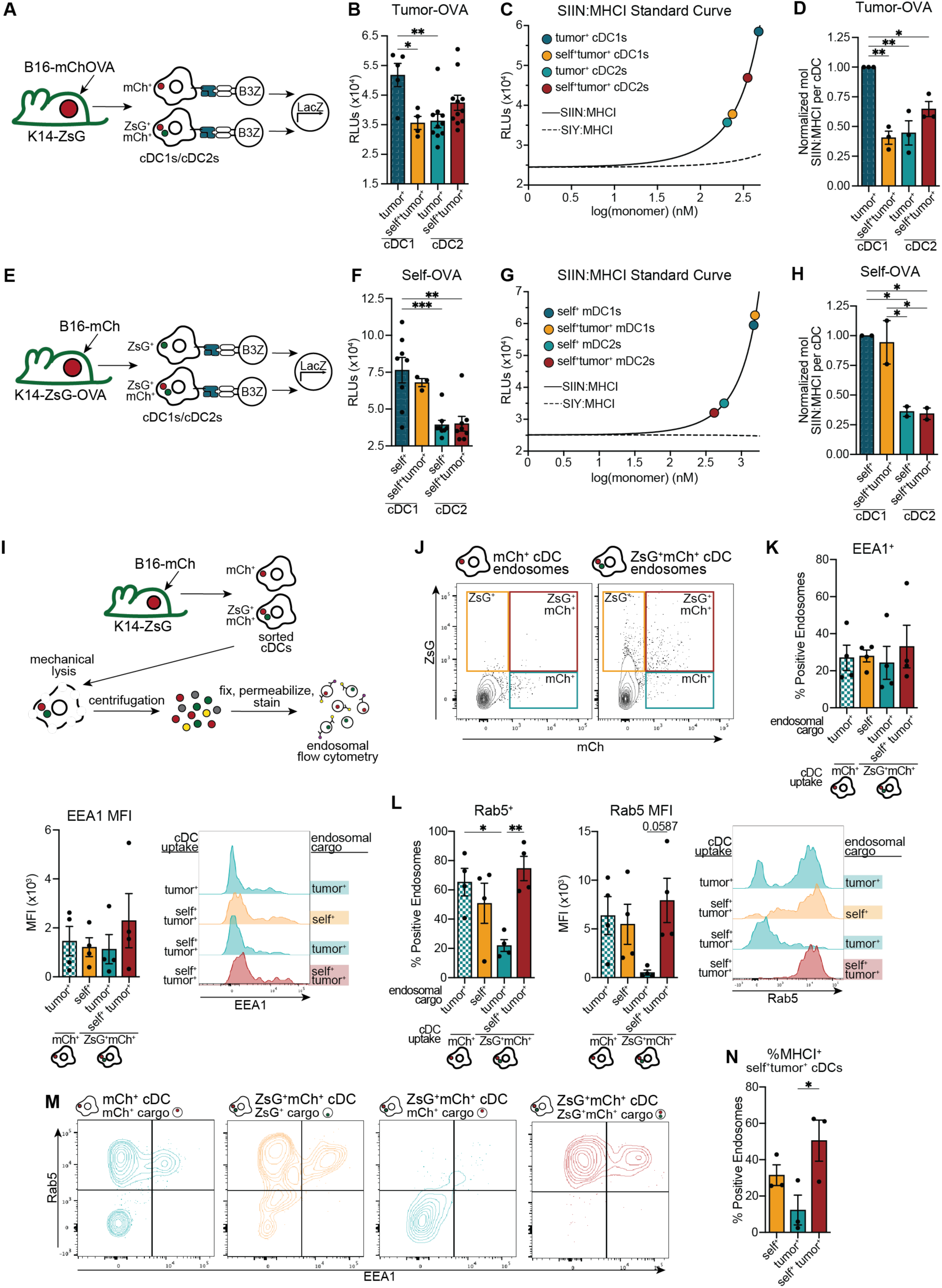
Concurrent uptake of self-antigen alters the source of peptide binding to surface MHCI in cDC1s. (A, E) Schematic of luminescent (A) tumor-specific or (E) self-specific B3Z assay with migratory cDC1s (mDC1s) and cDC2s (mDC2s) sorted by subset and either (A) tumor or (E) self uptake alongside self and tumor uptake. (B, F) RLUs of (B) tumor-specific or (F) self-specific B3Z coculture with antigen and subset sorted migratory cDCs plotted as raw counts. (C, G) Representative raw RLUs from (C) tumor-specific or (G) self-specific B3Z assay plotted against normalization curves with titrated SIIN:MHCI or SIY:MHCI monomers. (D, H) Fold change of normalized SIIN:MHCI quantification compared to monomer normalized mDC1 culture for (D) tumor-specific moles SIIN:MHCI or (H) self-specific moles SIIN:MHCI. (I) Schematic of endosomal phenotyping from intratumoral tumor^+^ or tumor^+^self^+^ cDCs. (J) Representative flow cytometry plots of endosomes isolated from tumor^+^ (left) or tumor^+^self^+^ (right) cDCs. (K-L) Percent positive (left), MFI (middle) and representative histogram (right) of (K) EEA1 and (L) Rab5 of endosomes from sorted cDCs. (M) Representative flow plots off EEA1 and Rab5 of gated on (far left) tumor^+^ endosomes of tumor^+^ cDCs, (near left) self+ endosomes, (near right) tumor^+^ endosomes, or (far right) self^+^tumor^+^ endosomes of self^+^tumor^+^ cDCs. (N) Percent MHCI^+^ endosomes from self^+^tumor^+^ sorted cDCs. (A-N) Data are shown as (B, F, K, L, N) individual values ± SEM or (D, H) mean ± SEM. (B, D, F, H, K, L, N) Statistical analysis was performed using one-way ANOVA, with Tukey’s multiple comparison test. 5-8 mice were pooled for each experiment. (B, D, F, H) n = 1-4 per condition per experiment or (K, L, N) n = 1 per condition per experiment, with (K, L) 4, (B, D, N) 3, or (F, H) 2 independent experiments. *p < 0.05, **p < 0.01, ***p < 0.001, ****p < 0.0001.

To determine whether the number of cDC self-peptide:MHCI was impacted by uptake of both tumor and self, we cocultured self^+^ or self^+^tumor^+^ cDC subsets sorted from tdLNs of K14-ZsG-OVA mice implanted with B16-mCh tumors (Fig. 6E). B3Z T cells stimulation was higher in cDC1 cocultures compared to cDC2 cocultures, regardless of uptake. Surprisingly, no differences were seen within the cDC1s based on antigen-load (Fig. 6F). Quantification of SIIN:MHCI using the standard curve supported this, revealing no significant differences in the number of SIIN:MHCI molecules on the surface of self^+^ cDC1s compared to self^+^tumor^+^ cDC1s (Fig. 6G-H). This suggested that, although self^+^tumor^+^ cDC1s promote more cross-priming of self-specific T cells compared to their self^+^ counterparts (Fig. 5N), the amount of self-antigen presented by self^+^ and self^+^tumor^+^ cDC1 is equivalent. Given no shift in total surface MHCI levels (Fig. 5K), this would indicate an overall decrease in the ratio of tumor:self antigen presented on cDC1s upon dual-antigen uptake.

Endosomal staining had revealed that antigen source could dictate the recruitment of cross presentation to antigen-bearing endosomes with *S. aur* driving robust MHCI enrichment in the EEA1+, Rab5+ cross-presenting endosomal compartment (Fig. 2F-H). However, the relationship between endosomal maturation, MHCI recruitment and antigen presentation in self^+^tumor^+^ cDCs was unknown. To determine whether cDC endosomal processing of self and tumor antigen was implicated in the decrease in tumor-peptide:MHCI on the surface of cDC1s, we sorted tumor^+^ cDCs and self^+^tumor^+^ cDCs from the TME and performed endosomal isolation and purification. We then stained the purified endosomes from each condition with endosomal markers EEA1 and Rab5 (Fig. 6I). This experiment would allow us to compare tumor^+^ endosomes from cDCs that have taken up only tumor and which we have shown drive robust T cell stimulation, to tumor^+^, self^+^tumor^+^ and self^+^ endosomes isolated from cDCs that have taken up both self and tumor which we have shown to have reduced cross-priming capability (Fig. 6J). Importantly, self^+^tumor^+^ cDCs contained a population of mCh^+^ZsG^+^ endosomes, indicating colocalization of tumor and self-antigen, consistent with our previous imaging analyses (Fig. 3H-I). We found no significant difference in EEA1 association with endosomes from either tumor^+^ or self^+^tumor^+^ cDCs, independent of endosomal cargo (Fig. 6K). In contrast, Rab5 showed specific association with tumor^+^ endosomes from tumor^+^ cDCs. This association was also seen in self^+^tumor^+^ cDCs, but was specific to the self^+^tumor^+^ endosomes, and not the tumor^+^ endosomes. This suggests that the tumor^+^ endosomes within the self^+^tumor^+^ cDCs fail to associate with the cross-presentation compartment, as marked by Rab5 (Fig. 6L-M). These data suggest that within self^+^tumor^+^ cDCs, endosomal processing for cross-presentation may favor self^+^tumor^+^ endosomes over tumor^+^ endosomes. Indeed, upon permeabilization and MHCI staining, we find that self^+^tumor^+^ endosomes have significantly more MHCI compared to tumor^+^ endosomes isolated from the same self^+^tumor^+^ cDCs (Fig. 6N). Processing and MHCI loading of antigens from self^+^tumor^+^ endosomes would be anticipated to result in both self and tumor-derived peptide loading onto MHCI. An apparent lack of processing and MHCI loading from tumor^+^ endosomes would lead to fewer peptides from these endosomes being presented. Therefore, preferential loading of self^+^tumor^+^ endosomes compared to tumor^+^ endosomes, but not self^+^ endosomes may align with the observed decrease in surface tumor-peptide:MHCI without the drop in surface self-peptide:MHCI seen in cDC1s (Fig. 6D, 6H). Endosomal colocalization and restriction of endosomal MHCI association may therefore serve as a novel mechanism involved in reducing tumor antigen cross-presentation by multiantigen bearing cDC1s.

## DISCUSSION

This study reveals a previously underappreciated capacity of cDCs to simultaneously sample, segregate, and process multiple antigen sources *in vivo*, and that this multiantigen handling has direct consequences for cross-presentation and CD8^+^ T cell priming. Using fluorescent antigen systems that allowed visualization of endosomal organization within individual cDCs, we found that cDCs can autonomously maintain distinct intracellular compartments for certain antigens (e.g. *S. aureus*:self, *S. aureus*:tumor), while permitting colocalization of other antigens (e.g. tumor:self). This organizational capacity becomes particularly consequential in the TME, where we observed that a substantial proportion of cDCs concurrently take up both tumor-derived and self-derived material. The functional outcome of this co-uptake is a striking shift in cross-presentation dynamics: cDC1s that have internalized both self and tumor antigen exhibit diminished tumor-specific CD8⁺ T cell priming, despite increased maturation, while simultaneously enhancing priming to self-derived antigen.

Using fluorescent model systems we identify and track non-cancerous ‘self’ (derived specifically from keratinocytes) in the melanoma TME. Within melanoma and other solid tumors, tumor-specific antigens that are the targets of antitumor immunity are expressed alongside various other non-cancerous ‘self’ derived antigens (28,52). These other sources of self are not tracked within our model system, but are likely still potential antigen sources across various solid tumor types capable of disrupting antitumor responses by cDCs. This would suggest the impact of cDC simultaneous processing of tumor and self likely extends beyond what we can detect using our tracking system, having widespread implications for tumor-specific T cell priming. Further, non-tumor antigen is not limited to host derived sources within the TME. Microbial (37,53) and fungal (54) communities can be present within solid tumors, adding an additional layer of complexity to the TME. In our study, we have identified that cDCs can access microbial derived antigen through unbroken skin and simultaneously process microbial and tumor derived antigen (Fig. 1O-1Q). We anticipate that PRR signaling for each of these antigen sources may distinctly impact how cDC sort and selectively process antigens for cross-presentation. Additional research into how cDCs handle various foreign and host antigen sources alongside tumor antigen may reveal strategies for improving tumor specific priming of T cells by cDCs.

Our findings both reinforce and complicate the prevailing paradigm that APCs autonomously process antigens based on antigen source specific cues. Consistent with prior *in vitro* work, we show that antigen processing by APCs is strongly influenced by antigen identity. Namely, only antigens associated with strong PRR, such as *S. aureus* efficiently recruit intraendosomal MHCI and show endosomal markers associated with cross-presentation (Fig. 2C-F) (12,14), However, our *in vivo* analyses show that cell derived self and tumor antigens remain endosomally segregated from *S. aureus* within the same cDC (Fig. 1E-1G, 1S-1U). This demonstrates that cDCs can maintain source specific compartmentalization even when multiple antigens are acquired.

The unexpected complexity arises when two non-microbial antigens, self and tumor, are co-internalized by an individual cDC. In this context, cDCs no longer maintain strict compartmental autonomy. Self and tumor antigens frequently colocalize within endosomes, creating a mixed compartment whose processing fate cannot be predicted from the behavior of either antigen alone. In this study, we determine that colocalization of self and tumor antigen leads to endosomal processing similar to tumor antigen from single antigen bearing cDCs, but appears to impact the processing of tumor only endosomes within the same multiantigen bearing cDCs (Fig. 6L-6M). In this way, colocalization of antigens can act as an antagonist to autonomous processing. Further *in vivo* studies that dissect endosomal processing dynamics will be essential to understand how antigen dependent PRR signaling, compartment colocalization and cross-presentation intersect to shape cDC function.

These findings have important implications for our understanding of antitumor immunity. First, they highlight that antigenic context within the TME is not merely background noise but an active determinant of cDC1 function. Recent work from Fan *et al.* further underscores how the origin and trafficking route of self-derived material can shape antigen presentation outcomes. They demonstrated that macrophages acquire cytosolic material from living cells through a trogocytosis-like mechanism that bypasses canonical degradative pathways and routes cargo into alternative vesicular compartments, resulting in altered CD8⁺ T cell priming (55). These findings highlight that self-derived cargo within APCs may be preferentially routed or preserved in ways that influence which peptides ultimately reach MHCI. In our studies, even when cDC1s acquire tumor antigen and upregulate classical maturation markers, the simultaneous presence of abundant tolerogenic self-antigen can blunt their ability to cross-prime tumor-specific CD8⁺ T cells. This helps explain why robust antigen uptake and maturation do not always translate into effective antitumor responses in vivo. Second, the effects we observe are subset-specific: cDC1s, but not cDC2s, exhibit impaired tumor cross-priming upon co-uptake of self-antigen, even though both subsets show similar levels of antigen uptake and maturation. This underscores that cDC1 functional specialization is not fixed but can be reshaped by the antigenic milieu. Third, these results suggest new therapeutic opportunities. Strategies that enhance tumor antigen routing into cross-presentation-competent compartments, stabilize tumor-derived peptides within those compartments, or reduce the competitive influence of self-antigen may restore or amplify cDC1-mediated antitumor immunity. Such approaches could complement existing immunotherapies.

In sum, this work identifies multiantigen processing within individual cDCs as a physiologically relevant feature of the TME that can reprogram endosomal processing and MHCI loading in ways that undermine tumor-specific CD8⁺ T cell priming. By uncovering how self-antigen uptake suppresses cDC1-mediated tumor cross-presentation while enhancing self-specific priming, our findings expose a previously unrecognized barrier to effective antitumor immunity. Targeting the intracellular decision-making processes that govern antigen routing and MHCI loading may offer new avenues to strengthen cDC1 function and improve immunotherapeutic outcomes.

### Limitations of the study

Several limitations should be considered. Our fluorescent antigen systems and use of model antigens such as OVA provide mechanistic clarity but do not capture the full diversity of endogenous tumor neoantigens or the complexity of human TMEs. The temporal resolution of our analyses provides snapshots of endosomal organization and peptide:MHCI abundance but does not yet define the dynamic evolution of these processes over time. Additionally, our work focuses on skin-associated tumors and keratinocyte-derived self-antigen; other tissues may present different balances of antigen sources and distinct patterns of cDC subset behavior. Nonetheless, the principles uncovered here, particularly the influence of antigenic competition on endosomal routing and MHCI loading, are likely to be broadly relevant across tissues.

## Acknowledgments

We thank M. Krummel for providing the K14-ZsGreen mice and B16F10 reporter cell lines. We thank E. Alspach for discussion and support. Funding: This work was supported by Harry J. Lloyd Charitable Trust (M.K.R.), the V Foundation V Scholar Award (M.K.R), the Concern Foundation Conquer Cancer Now Award (M.K.R.), NCI R01CA299335 (M.K.R.), T32AI170496 (V.P.S.), T32GM142619 (K.B.), NIH R01CA245535 (N.O), American Cancer Society RSG-24-1248754-01-MM (N.O.), NIAMS R01AR080034 (T.C.S). We acknowledge the OHSU Flow Cytometry Shared Resource (RRID SCR_009974) for assistance generating flow cytometry data. Research reported here was supported in part by the Knight Cancer Institute CCSG grant NIH P30CA069533.

## MATERIALS AND METHODS

### Animal Models

Mice were housed in the specific pathogen-free environment at the Oregon Health and Science University (OHSU) mouse facility at the South Waterfront campus in Portland, OR. All experimental protocols were preapproved by The Institutional Animal Care and Use Committee (IACUC) at OHSU under parent protocol number IP00003351. Mouse procedures were performed by approved personnel, and care was provided by OHSU’s Department of Comparative Medicine. Mice used for the study were received from indicated sources and then bred and maintained in house on a C57BL/6 background:

Wildtype: C57BL/6, strain #000664 purchased from The Jackson Laboratory.

OTI: C57BL/6-Tg(TcraTcrb)1100Mjb/J, strain #003831 purchased from The Jackson Laboratory.

K14-Cre; lox-stop-lox-ZsGreen: Gifted from M. Krummel (39). Generated by crossing K14-Cre (B6N.Cg-Tg(KRT14-cre)1Amc/J, strain #018964 Jackson Labs) with Ai6 strain (B6.Cg-Gt(ROSA)26Sor^tm6(CAG-ZsGreen1)Hze^/J, strain #007906).

K14-Cre; lox-stop-lox-ZsGreen; K14-OVA: Generated by crossing K14-Cre; lox-stop-lox-ZsGreen strain with K14-OVA (C57BL/6-Tg(KRT14-OVAL*)15Sika/J, strain #026562 originally purchased from Jackson labs) (44).

K14-Cre; lox-stop-lox-tdTomato: Generated by crossing K14-Cre^+^; Ai6^-/-^ strain with Ai14 strain (B6.Cg-Gt(ROSA)26Sor^tm14(CAG-tdTomato)Hze^/J, strain #007914 Jackson Labs) gifted from J. Moreau, as shown in Fig 3.S1A and draining lymph node specific trafficking of keratinocyte derived antigen by cDCs verified in Fig. 3.S1B.

A combination of 6- to 16-week-old male and female mice was used for all experiments. Pooled experiments included a combination of both sexes. Mice were randomly assigned to experimental groups whenever possible. Same-sex littermates were housed together in the same cages. Genotypes were confirmed by in-house PCR analysis. All mice were kept on a 12-hour light/12-hour dark cycle. For tumor studies, mice were monitored for pain and distress and euthanized before tumors reached 2cm in diameter. Adult mice were euthanized at defined endpoints with CO2, followed by secondary cervical dislocation, and neonates were euthanized via decapitation.

### Cell lines

B16-ZsGreen, B16-ZsG-OVA, B16-mCherry and B16-mChOVA cell lines were generated and validated as described previously (39,56). In brief, the B16-F10 cell line (ATCC, CRL-6475) was virally transduced with constructs for ZsGreen, ZsGreen-OVA, mCherry, or mCherry-OVA. Following transduction, cells were FACs-sorted for stable ZsGreen or mCherry expression. Tumor cell lines were cultured with D10 media (DMEM (Gibco), 10% heat-inactivated FBS (HyClone), and 1% Penicillin-Streptomycin-Glutamine (Gibco)).

The B3Z cell line has been described previously (57) and was a gift from M. Gough. The cell line was maintained in R10 (RPMI1640 (Gibco), 10% heat-inactivated FBS (HyClone), and 1% Penicillin-Streptomycin-Glutamine (Gibco)) supplemented with 1% HEPES solution (MilliporeSigma), 1% sodium pyruvate solution (Sigma-Aldrich), 1% MEM-NEAA (Fisher Scientific), and 0.1% 2-Mercaptoethanol (Fisher Scientific).

Keratinocyte lineage derivation has been described previously (38). Neonatal (P0 to P1) mice of K14-ZsGreen and K14-ZsGreen-OVA were removed from the mother’s cage and kept isolated to prevent contamination. Neonates were euthanized by rapid decapitation. Tail samples were taken to confirm genotype of K14-Cre+; Ai6+/- and K14-Cre+; Ai6+/-;K14-OVA+. Dorsal skin was carefully removed from the mice and digested with 1 mL of Dispase II (Aldrich) at 37°C for 2.5 hours. Epidermis was carefully separated from the dermal layer with forceps and spread into 1 mL of TrypLE (Gibco). Epidermal sheets were incubated at 24°C for 10 minutes. E50 medium (defined previously (38) as “E Media”, gifted from N. Oshimori) was pipetted into each sample, and basal keratinocytes were gently pipetted to detach from the epidermal layer. Samples were collected, centrifuged (400 x g, 3 min), and washed with 1 mL of E50 medium. The samples were then transferred into a 6-well plate containing mitomycin C (Cayman Chemical Company, 10 μg/mL) treated 3T3 feeder cells (gifted from N. Oshimori) at approximately 50% confluence in 2.5 mL of E50 medium. The cultures were maintained for at least 10 passages in E50 medium prior to freezing stocks, and fluorescence was confirmed through fluorescence microscopy and flow cytometry, as shown in Fig. 3.S2A.

All lines for this study were maintained at 5% CO2, 37°C, and 95% humidity. All cell lines were routinely tested for Mycoplasma contamination. Cell line identities were regularly confirmed through fluorescent reporter expression and flow cytometry.

### BMDC Generation

Femurs and tibiae from wildtype mice were isolated and cleaned of tissue. The heads of the bones were trimmed, and bone marrow was collected by flash centrifugation into 1.5 mL microcentrifuge tubes. Cells were resuspended in 3 mL of ACK lysis buffer (Thermo Fisher Scientific) and incubated at 24 °C for 3 minutes to lyse red blood cells. D10 medium was added to quench lysis, and cells were pelleted (400 × g, 5 min), filtered through a 70 μm cell strainer (Corning), and resuspended to 1 x 10⁶ cells/mL in D10 supplemented with 0.1% 2-mercaptoethanol (Fisher Scientific), 3.75 ng/mL GM-CSF (PeproTech), and 100 ng/mL FLT3L (Bio X Cell). Cells were plated at 10 mL per well in 6-well plates (Corning). BMDCs were maintained at 37°C with 5% CO₂ and 95% humidity for 7–14 days post-harvest. On days 3-4, BMDC cultures were supplemented with fresh 3.75 ng/mL GM-CSF and 100 ng/mL FLT3L. BMDCs were replated on day 7 in fresh D10 medium supplemented with 0.1% 2-mercaptoethanol, 3.75 ng/mL GM-CSF, and 100 ng/mL FLT3L, and maintained until harvest for experiments (no later than 14 days post-harvest).

### Tumor Injections

B16-ZsGreen, B16-ZsG-OVA, B16-mCherry, and B16-mChOVA tumor cell lines were maintained in culture until approximately 80% confluency. Cells were washed with sterile DPBS (Corning), detached using 0.05% Trypsin-EDTA (Corning), washed with sterile DPBS, and resuspended to 10^7^ cells/mL in RPMI1640 (Gibco). Resuspended cells were combined with an equal volume of Matrigel matrix (Corning) for a 1:1 (v/v) mixture for injections. Mice were injected subcutaneously with 50 μL bilateral injections into shaved dorsal flanks (2.5 x 10^5^ cells/injection). Tumors and draining lymph nodes were harvested at defined experimental time points or when tumors reached IACUC-approved humane endpoints.

### Microbe Prep and Association

Fluorescent *Staphylococcus aureus* (S. *aur*) expressing ZsGreen (S. *aur*-ZsG) and S. *aur* expressing mCherry (S. *aur*-mCh) were generated from USA300 strains SF8300 with constructs pJL74-2w-zsgreen and pJL74-CAM-2w-mcherry, respectively, as described previously (36) (gifted as glycerol stocks from T. Scharschmidt). Glycerol stocks of the strains were streaked out onto tryptic soy broth (TSB, BD Bacto) agar (Fisher Sci.) plates containing erythromycin (10 μg/mL, Fisher Sci.) or chloramphenicol (15 μg/mL, Acros Organics) and cultured overnight at 37°C. The following day, single colonies were picked and used to inoculate liquid TSB supplemented with selective antibiotics and cultured in shaker overnight, 180rpm, 32°C. In the morning, cultures were collected, measured for OD600 to determine concentration, washed with sterile PBS, and centrifuged (10 min, 4°C, 3000rpm) to resuspend to 1 x 10^9^ CFUs/mL in sterile PBS on ice. For cutaneous applications, 200 μL of resuspended microbe was ‘painted’ onto either to the dorsal skin of shaved mice or 100 μL was painted atop each bilateral tumor. Tissues with microbes were harvested after approximately 16 hours after skin or tumor association.

### Lymph Node Digest

Skin- and tumor- draining inguinal, axillary, and brachial lymph nodes (dLNs) were collected from mice as previously described (56,39). The dLNs were transferred into 24-well plates (Corning) with cold RPMI1640 medium (Gibco). Digestion media was prepped: RPMI1640 containing 100 U/mL Collagenase Type I (Worthington Biochemical), 500 U/mL Collagenase Type IV (Worthington Biochemical), and 20 μg/mL DNase I (Roche). A total of 1 mL digestion media was used for each set of 6 pooled dLNs from an individual mouse. The dLN capsules were dissected using fine forceps and incubated in digestion medium at 37°C for 30 minutes, pipetting to disrupt after 15 minutes. 1 mL of MACS buffer (PBS (Gibco), 2% heat-inactivated FBS (HyClone), 1 mM EDTA (Fisher BioReagents), 1% Penicillin-Streptomycin-Glutamine (Gibco), and 2 mM HEPES (MilliporeSigma) was added to each sample. Digested dLN samples were pooled if appropriate and suspensions were passed through 70 μm strainers (Corning). Samples were centrifuged (400 × g, 5 min). For sorting experiments, samples were enriched with the EasySep™ Mouse Pan-DC Enrichment Kit II (STEMCELL Technologies) and magnet per manufacturer’s recommendations prior to staining.

### Cell Staining

Collected cells were stained with Zombie viability dye (BioLegend) diluted in DPBS for 15 min on ice. Cells were washed and stained for surface markers with Fc-block (BD Biosciences) in MACS buffer for 20 min on ice. Samples were washed with MACS buffer, centrifuged (400 × g, 5 min), and resuspended in MACs (or DAPI diluted in MACS if Zombie not used) for flow cytometry or FACS sorting.

For intracellular staining, cells were pelleted (400 × g, 5 min) and resuspended in R10 media: RPMI1640 (Gibco), 10% heat-inactivated FBS (HyClone), 1% Penicillin-Streptomycin-Glutamine (Gibco), and 0.1% 2-mercaptoethanol (Gibco). GolgiPlug (BD Biosciences) was added at 1 μL/mL and cells were incubated for 5 h at 37°C prior to proceeding with Zombie staining. After washing, Zombie dye, and surface staining as described above, cells were fixed and permeabilized using Cytofix/Cytoperm solution (BD Biosciences) for 20 min on ice. Then samples were stained with intracellular antibody staining diluted in Cytofix/Cytoperm wash buffer with Fc-block for 1 hour on ice. Each antibody had a corresponding fluorescence minus one (FMO) control for gating. Samples were washed with Cytofix/Cytoperm wash buffer, centrifuged (400 × g, 5 min), and resuspended in MACS for flow cytometry analysis.

### Flow Cytometry and Sorting

Cell sorting was performed using a BD FACSymphony S6 (BD Biosciences) equipped with a 100 μm nozzle with BD FACSDiva software (BD Biosciences, v9). Samples were sorted as a four-way sort into 1.5 mL Eppendorf tubes containing 700 μL D10 supplemented with 0.1% 2-mercaptoethanol and 3.75 ng/mL GM-CSF to maintain viability during extended sorts. Tubes were kept protected from light and maintained at 4°C for the sort duration. For live cell imaging experiments, DMEM in D10 was replaced with phenol red-free RPMI1640 (Gibco). Following collection, sorted cells were centrifuged (400 × g, 5 min) and resuspended in appropriate media for downstream applications. Flow cytometry acquisition was performed on a BD FACSymphony A5 (BD Biosciences), and data were analyzed using FlowJo software (Tree Star, v10).

### Tumor and Skin Tissue Digest

Back skin or tumors were harvested from mice, collected in RPMI1640, and mechanically dissociated with scissors or razor blades prior to digestion. For skin digestion, minced tissue was incubated in D10 containing collagenase XI (Sigma-Aldrich), hyaluronidase (Sigma-Aldrich), and DNase I (Sigma-Aldrich), supplemented with cytochalasin D (Cayman Chemical) and gentamicin (Acros Organics, if *S. aur* used in model). Skin samples were digested in approximately 4 mL per skin sample at 37°C with shaking (200 rpm) for 50 min. Collected tumors were digested in RPMI1640 with Collagenase Type I (Worthington Biochemical), Collagenase Type IV (Worthington Biochemical), and DNase I (Roche). Tumor samples were digested in approximately 3 mL per tumor at 37°C with shaking (200 rpm) for 40 min.

Following digestion, samples were diluted with D10, pooled if appropriate for the experiment, filtered through 100 μm strainers (Corning), centrifuged (400 × g, 5 min), and resuspended in cold MACS buffer. For experiments requiring cell sorting, CD45^+^ immune cells were positively enriched for using the EasySep Mouse CD45 Positive Selection Kit (STEMCELL Technologies) according to the manufacturer’s instructions prior to antibody staining. All samples were subsequently stained with antibody panels as described.

### OT-I T Cell Stimulation Assays

Spleens were harvested from transgenic OT-I mice and mechanically dissociated using a plunger through a 70 μm strainer into single-cell suspensions. Splenocyte preparations were pelleted (400 x g, 5 min) and resuspended in 3 mL ACK lysis buffer (Gibco) for a 3 min, 24°C incubation to lyse red blood cells prior to dilution in MACS buffer. Samples were centrifuged (400 × g, 5 min), resuspended in MACS, filtered through 70 μm strainers, and enriched for CD8^+^ T cells using the negative selection CD8^+^ T cell isolation kit (STEMCELL Technologies) per manufacturer’s instructions. Isolated CD8^+^ T cells were washed and labeled with 1 μM Violet Proliferation Dye (BD Biosciences) diluted in DPBS for 15 min at 37°C. Cells were then washed, resuspended in R10 media at 5 × 10^5^ cells/mL (bulk cDCs) or 2.5 × 10^5^ cells/mL (cDC subsets). Sorted cDC populations were separately resuspended at 5 × 10^4^ cells/mL (bulk) or 2.5 × 10^4^ cells/mL (cDC subsets). Cocultures were established in 96-well V-bottom plates at a 10:1 T cell:cDC ratio by combining equal volumes of each cell suspension for a total volume of 200 μL per well. Cultures were maintained at 37°C with 5% CO₂ and 95% humidity for 72 hours prior to flow cytometric analysis. T cell activation and proliferation were evaluated using the gating strategy shown in Figure 2A.

### Imaging and Colocalization Analysis

Sorted cells were resuspended in phenol red-free R10 media and plated onto fibronectin-coated 384-well glass-bottom plates containing high-performance #1.5 cover glass (CellVis) at approximately 2.5 × 10^3^ cells in 100 μL per well. Cells were allowed to adhere for a minimum of 1 h in 37°C, 5% CO₂, humidified incubator prior to imaging.

Live-cell fluorescence imaging was performed using a Yokogawa CSU-X1 spinning disk confocal system mounted on a Zeiss Axio Observer microscope equipped with a Plan-Apochromat 63×/1.40 NA oil immersion DIC M27 objective. Samples were maintained at 37°C with 5% CO₂ in a humidified environmental chamber throughout image acquisition. Images were acquired using ZEN Blue software (Zeiss, v2.6) with sequential acquisition of green then red fluorescence channels with a single intermediate brightfield plane within the middle of the Z-stack. Z-stack images were acquired at 0.27 μm Z-step intervals throughout the imaged cell volume.

Image processing and colocalization analyses were performed using Fiji/ImageJ with the DiAna plugin (58). Analyses were performed within individual experimental conditions using identical threshold and analysis parameters to maintain consistency within individual experimental conditions. Three-dimensional colocalization and distance analyses were performed on reconstructed image stacks to assess antigen localization within individual cDCs. Colocalization measurements were normalized to the limiting quantity of antigens within each analyzed cell. Cells in direct physical contact through dendritic processes were excluded from analysis to reduce antigen transfer. Non-colocalized antigens were calculated as 100% minus the colocalized fraction within each individual cell. Representative images were adjusted individually for threshold intensity and generated solely for visualization purposes.

### BMDC Cocultures

BMDCs were generated as described above and cocultured with fluorescent *S. aureus* strains and/or mitomycin C-treated (Cayman Chemical) K14-ZsGreen-OVA or B16-ZsGreen-OVA cells. K14-ZsGreen-OVA and B16-ZsGreen-OVA cultures were treated with 20 μg/mL or 25 μg/mL mitomycin C, respectively, for 21–23 h prior to coculture harvest to induce apoptotic cell death. Supernatants containing apoptotic material were retained during cell harvest and recombined with collected cells for preparation. Fluorescent *S. aureus* strains were cultured as described above and heat-killed at 70°C for 90 min prior to coculture to prevent active infection. Heat-killed bacteria were washed twice with DPBS and pelleted by centrifugation (3000 rpm, 10 min, 4°C). Successful heat killing was confirmed by streaking treated cultures onto selective agar plates. BMDCs, apoptotic cell preparations, and heat-killed bacteria were resuspended in antibiotic-free D10 media and combined at a ratio of 1:1:10 (BMDCs:B16/K14 cells:*S. aureus* MOI). Cocultures were plated in 96-well plates at a final volume of 200 μL per well and maintained at 37°C with 5% CO₂ for 4 h with mixing every 1 hour. Cocultures were then either harvested for flow cytometry to phenotype cells or processed for vesicle flow cytometry.

### Vesicle Flow Cytometry

Vesicle flow cytometry was adapted and optimized from prior protocols (40). Following coculture, BMDCs were harvested and CD11c^+^ cells were enriched using a biotinylated anti-CD11c antibody together with magnetic bead enrichment reagents (STEMCELL Technologies) according to established protocols (39). Enriched cells or sorted cDCs were washed in cold PBS and resuspended in homogenization buffer: 8% sucrose (Fisher Sci.), 3mM imidazole (Sigma-Aldrich), 1mM DTT (Fisher Sci.), 1 tablet cOmplete, Mini, EDTA-free Protease Inhibitor Cocktail (Roche) in 10mL DPBS. Samples were pushed through a 22 x g needle 30 times. Post-nuclear supernatants were isolated by centrifugation (150 × g, 4 min, 4°C), transferred to fresh wells, and vesicle-containing fractions were collected by centrifugation at 3000 x rpm, 4 min, 4°C. Vesicle pellets were washed in cold PBS prior to downstream surface and ICS cell staining, as described. Vesicle samples were analyzed on a BD FACSymphony A5 (BD Biosciences). Size calibration beads (Spherotech, Inc.) were included during acquisition to assist vesicle gating and population identification. Data analysis was performed using FlowJo software (Tree Star, v10).

### B3Z T Cell Stimulation Assays

Sorted cDC subsets were resuspended to 1.8 × 10^3^ cDCs / 25 μL and B3Z cells were resuspended to 1.8 × 10^2^ B3Z cells / 25 μL in B3Z assay media: phenol red-free R10 media supplemented with 1% MEM non-essential amino acids (Fisher Sci.), 1% sodium pyruvate (Sigma-Aldrich), 1% HEPES (Sigma-Aldrich), and 0.1% 2-mercaptoethanol. Equal volumes of cells (25 μL) were added to each well for a 10:1, cDC:B3Z ratio coculture. Assays were performed in flat-bottom, white tissue culture-treated 384-well plates with a final volume of 50 μL per well consisting of 25 μL B3Z suspension combined with 25 μL cDC suspension or peptide-loaded monomer controls. For monomer titration controls, H-2Kb:SIINFEKL or H-2Kb:SIYRYYGL monomers (NIH Tetramer Core Facility) were serially diluted in B3Z assay media and added in place of cDCs. H-2Kb:SIINFEKL monomers were diluted 1:2 from 500 nM to 0.244 nM, while H-2Kb:SIYRYYGL monomers were diluted 1:4 from 500 nM to 0.122 nM. Cocultures were incubated for 18 hours at 37°C with 5% CO₂ in a humidified incubator. Following incubation, 25 μL Beta-Glo Luciferase Assay substrate (Promega) was added to each well and plates were incubated for 30 min at room temperature protected from light prior to luminescence acquisition using a plate reader with 1 sec integration time per well. Experimental cDC RLUs were compared against nonlinear fit curves with bottom constraints equal to mean of triplicate wells with B3Z cells with B3Z assay media only.

## REFERENCES

1. Carroll SL, Pasare C, Barton GM. Control of adaptive immunity by pattern recognition receptors. Immunity. Elsevier; 2024;57:632–48.

2. Ohara RA, Murphy KM. Recent progress in type 1 classical dendritic cell cross-presentation - cytosolic, vacuolar, or both? Curr Opin Immunol. 2023;83:102350.

3. Yee Mon KJ, Blander JM. TAP-ing into the cross-presentation secrets of dendritic cells. Curr Opin Immunol. 2023;83:102327.

4. Huang AYC, Bruce AT, Pardoll DM, Levitsky HI. In Vivo Cross-Priming of MHC Class I–Restricted Antigens Requires the TAP Transporter. Immunity. 1996;4:349–55.

5. Bertholet S, Goldszmid R, Morrot A, Debrabant A, Afrin F, Collazo-Custodio C, et al. *Leishmania* Antigens Are Presented to CD8+ T Cells by a Transporter Associated with Antigen Processing-Independent Pathway In Vitro and In Vivo. J Immunol. 2006;177:3525–33.

6. Gros M, Segura E, Rookhuizen DC, Baudon B, Heurtebise-Chrétien S, Burgdorf N, et al. Endocytic membrane repair by ESCRT-III controls antigen export to the cytosol during antigen cross-presentation. Cell Rep. 2022;40:111205.

7. Barbet G, Nair-Gupta P, Schotsaert M, Yeung ST, Moretti J, Seyffer F, et al. TAP dysfunction in dendritic cells enables noncanonical cross-presentation for T cell priming. Nat Immunol. Nature Publishing Group; 2021;22:497–509.

8. Sengupta D, Graham M, Liu X, Cresswell P. Proteasomal degradation within endocytic organelles mediates antigen cross-presentation. EMBO J. 2019;38:EMBJ201899266.

9. Cebrian I, Visentin G, Blanchard N, Jouve M, Bobard A, Moita C, et al. Sec22b Regulates Phagosomal Maturation and Antigen Crosspresentation by Dendritic Cells. Cell. Elsevier; 2011;147:1355–68.

10. Guermonprez P, Saveanu L, Kleijmeer M, Davoust J, van Endert P, Amigorena S. ER–phagosome fusion defines an MHC class I cross-presentation compartment in dendritic cells. Nature. 2003;425:397–402.

11. Blander JM, Medzhitov R. Regulation of Phagosome Maturation by Signals from Toll-Like Receptors. Science. 2004;304:1014–8.

12. Blander JM, Medzhitov R. Toll-dependent selection of microbial antigens for presentation by dendritic cells. Nature. Nature Publishing Group; 2006;440:808–12.

13. Burgdorf S, Schölz C, Kautz A, Tampé R, Kurts C. Spatial and mechanistic separation of cross-presentation and endogenous antigen presentation. Nat Immunol. Nature Publishing Group; 2008;9:558–66.

14. Hoffmann E, Kotsias F, Visentin G, Bruhns P, Savina A, Amigorena S. Autonomous phagosomal degradation and antigen presentation in dendritic cells. Proc Natl Acad Sci. 2012;109:14556–61.

15. Alloatti A, Kotsias F, Pauwels A-M, Carpier J-M, Jouve M, Timmerman E, et al. Toll-like Receptor 4 Engagement on Dendritic Cells Restrains Phago-Lysosome Fusion and Promotes Cross-Presentation of Antigens. Immunity. Elsevier; 2015;43:1087–100.

16. Schulz O, Diebold SS, Chen M, Näslund TI, Nolte MA, Alexopoulou L, et al. Toll-like receptor 3 promotes cross-priming to virus-infected cells. Nature. Nature Publishing Group; 2005;433:887–92.

17. Fang H, Ang B, Xu X, Huang X, Wu Y, Sun Y, et al. TLR4 is essential for dendritic cell activation and anti-tumor T-cell response enhancement by DAMPs released from chemically stressed cancer cells. Cell Mol Immunol. 2014;11:150–9.

18. Pirillo C, Al Khalidi S, Sims A, Devlin R, Zhao H, Pinto R, et al. Co-transfer of antigen and contextual information harmonises peripheral and lymph node cDC activation. Sci Immunol. 2023;8:eadg8249.

19. Chen DS, Mellman I. Oncology Meets Immunology: The Cancer-Immunity Cycle. Immunity. 2013;39:1–10.

20. Mellman I, Chen DS, Powles T, Turley SJ. The cancer-immunity cycle: Indication, genotype, and immunotype. Immunity. 2023;56:2188–205.

21. Hildner K, Edelson BT, Purtha WE, Diamond M, Matsushita H, Kohyama M, et al. Batf3 deficiency reveals a critical role for CD8alpha+ dendritic cells in cytotoxic T cell immunity. Science. New York, N.Y.; 2008;322:1097–100.

22. Böttcher JP, Bonavita E, Chakravarty P, Blees H, Cabeza-Cabrerizo M, Sammicheli S, et al. NK Cells Stimulate Recruitment of cDC1 into the Tumor Microenvironment Promoting Cancer Immune Control. Cell. 2018;172:1022–1037.e14.

23. Ferris ST, Durai V, Wu R, Theisen DJ, Ward JP, Bern MD, et al. cDC1 prime and are licensed by CD4+ T cells to induce anti-tumour immunity. Nature. Nature Publishing Group; 2020;584:624–9.

24. Salmon H, Idoyaga J, Rahman A, Leboeuf M, Remark R, Jordan S, et al. Expansion and Activation of CD103+ Dendritic Cell Progenitors at the Tumor Site Enhances Tumor Responses to Therapeutic PD-L1 and BRAF Inhibition. Immunity. 2016;44:924–38.

25. Roberts EW, Broz ML, Binnewies M, Headley MB, Nelson AE, Wolf DM, et al. Critical Role for CD103+/CD141+ Dendritic Cells Bearing CCR7 for Tumor Antigen Trafficking and Priming of T Cell Immunity in Melanoma. Cancer Cell. 2016;30:324–36.

26. Galeano Niño JL, Wu H, LaCourse KD, Kempchinsky AG, Baryiames A, Barber B, et al. Effect of the intratumoral microbiota on spatial and cellular heterogeneity in cancer. Nature. Nature Publishing Group; 2022;611:810–7.

27. Kalaora S, Nagler A, Nejman D, Alon M, Barbolin C, Barnea E, et al. Identification of bacteria-derived HLA-bound peptides in melanoma. Nature. Nature Publishing Group; 2021;592:138–43.

28. Nirmal AJ, Maliga Z, Vallius T, Quattrochi B, Chen AA, Jacobson CA, et al. The Spatial Landscape of Progression and Immunoediting in Primary Melanoma at Single-Cell Resolution. Cancer Discov. 2022;12:1518–41.

29. Canton J, Blees H, Henry CM, Buck MD, Schulz O, Rogers NC, et al. The receptor DNGR-1 signals for phagosomal rupture to promote cross-presentation of dead cell-associated antigens. Nat Immunol. 2021;22:140–53.

30. Caronni N, Piperno GM, Simoncello F, Romano O, Vodret S, Yanagihashi Y, et al. TIM4 expression by dendritic cells mediates uptake of tumor-associated antigens and anti-tumor responses. Nat Commun. Nature Publishing Group; 2021;12:2237.

31. Woo S-R, Fuertes MB, Corrales L, Spranger S, Furdyna MJ, Leung MYK, et al. STING-Dependent Cytosolic DNA Sensing Mediates Innate Immune Recognition of Immunogenic Tumors. Immunity. 2014;41:830–42.

32. Belz GT, Behrens GMN, Smith CM, Miller JFAP, Jones C, Lejon K, et al. The CD8α+ Dendritic Cell Is Responsible for Inducing Peripheral Self-Tolerance to Tissue-associated Antigens. J Exp Med. 2002;196:1099–104.

33. Waithman J, Allan RS, Kosaka H, Azukizawa H, Shortman K, Lutz MB, et al. Skin-Derived Dendritic Cells Can Mediate Deletional Tolerance of Class I-Restricted Self-Reactive T Cells1. J Immunol. 2007;179:4535–41.

34. Bedoui S, Whitney PG, Waithman J, Eidsmo L, Wakim L, Caminschi I, et al. Cross-presentation of viral and self antigens by skin-derived CD103+ dendritic cells. Nat Immunol. 2009;10:488–95.

35. Igyártá BZ, Haley K, Ortner D, Bobr A, Gerami-Nejad M, Edelson BT, et al. Skin-Resident Murine Dendritic Cell Subsets Promote Distinct and Opposing Antigen-specific T Helper Responses. Immunity. 2011;35:260–72.

36. Leech JM, Dhariwala MO, Lowe MM, Chu K, Merana GR, Cornuot C, et al. Toxin-Triggered Interleukin-1 Receptor Signaling Enables Early-Life Discrimination of Pathogenic versus Commensal Skin Bacteria. Cell Host Microbe. 2019;26:795–809.e5.

37. Nejman D, Livyatan I, Fuks G, Gavert N, Zwang Y, Geller LT, et al. The human tumor microbiome is composed of tumor type–specific intracellular bacteria. Science. American Association for the Advancement of Science; 2020;368:973–80.

38. Nowak JA, Fuchs E. Isolation and Culture of Epithelial Stem Cells. Methods Mol Biol Clifton NJ. 2009;482:215–32.

39. Ruhland MK, Roberts EW, Cai E, Mujal AM, Marchuk K, Beppler C, et al. Visualizing Synaptic Transfer of Tumor Antigens among Dendritic Cells. Cancer Cell. 2020;37:786–799.e5.

40. Savina A, Vargas P, Guermonprez P, Lennon A-M, Amigorena S. Measuring pH, ROS Production, Maturation, and Degradation in Dendritic Cell Phagosomes Using Cytofluorometry-Based Assays. In: Naik SH, editor. Dendritic Cell Protoc [Internet]. Totowa, NJ: Humana Press; 2010 [cited 2022 Apr 25]. page 383–402. Available from: http://link.springer.com/10.1007/978-1-60761-421-0_25

41. Huynh KK, Eskelinen E-L, Scott CC, Malevanets A, Saftig P, Grinstein S. LAMP proteins are required for fusion of lysosomes with phagosomes. EMBO J. 2007;26:313–24.

42. Burgdorf S, Kautz A, Böhnert V, Knolle PA, Kurts C. Distinct Pathways of Antigen Uptake and Intracellular Routing in CD4 and CD8 T Cell Activation. Science. American Association for the Advancement of Science; 2007;316:612–6.

43. Wang YY, Hu CF, Li J, You X, Gao FG. Increased translocation of antigens to endosomes and TLR4 mediated endosomal recruitment of TAP contribute to nicotine augmented cross-presentation. Oncotarget. Impact Journals; 2016;7:38451–66.

44. Schuster VP, Brown K, Unsworth JR, Berkowitz N, Ruhland MK. Contextual control of CD8+ T cell priming by dendritic cell subsets in tumor and inflammatory microenvironments [Internet]. bioRxiv; 2025 [cited 2026 Mar 31]. page 2025.10.31.685861. Available from: https://www.biorxiv.org/content/10.1101/2025.10.31.685861v1

45. Broz ML, Binnewies M, Boldajipour B, Nelson AE, Pollack JL, Erle DJ, et al. Dissecting the tumor myeloid compartment reveals rare activating antigen-presenting cells critical for T cell immunity. Cancer Cell. 2014;26:638–52.

46. Hubo M, Trinschek B, Kryczanowsky F, Tuettenberg A, Steinbrink K, Jonuleit H. Costimulatory Molecules on Immunogenic Versus Tolerogenic Human Dendritic Cells. Front Immunol. 2013;4:82.

47. Fonteneau JF, Kavanagh DG, Lirvall M, Sanders C, Cover TL, Bhardwaj N, et al. Characterization of the MHC class I cross-presentation pathway for cell-associated antigens by human dendritic cells. Blood. 2003;102:4448–55.

48. Lizée G, Basha G, Tiong J, Julien J-P, Tian M, Biron KE, et al. Control of dendritic cell cross-presentation by the major histocompatibility complex class I cytoplasmic domain. Nat Immunol. Nature Publishing Group; 2003;4:1065–73.

49. Orabona C, Grohmann U, Belladonna ML, Fallarino F, Vacca C, Bianchi R, et al. CD28 induces immunostimulatory signals in dendritic cells via CD80 and CD86. Nat Immunol. Nature Publishing Group; 2004;5:1134–42.

50. Keller AM, Schildknecht A, Xiao Y, van den Broek M, Borst J. Expression of Costimulatory Ligand CD70 on Steady-State Dendritic Cells Breaks CD8+ T Cell Tolerance and Permits Effective Immunity. Immunity. 2008;29:934–46.

51. Karttunen J, Sanderson S, Shastri N. Detection of rare antigen-presenting cells by the lacZ T-cell activation assay suggests an expression cloning strategy for T-cell antigens. Proc Natl Acad Sci. Proceedings of the National Academy of Sciences; 1992;89:6020–4.

52. Liu S, Liu C, He Y, Li J. Benign non-immune cells in tumor microenvironment. Front Immunol. 2025;16:1561577.

53. Galeano Niño JL, Wu H, LaCourse KD, Kempchinsky AG, Baryiames A, Barber B, et al. Effect of the intratumoral microbiota on spatial and cellular heterogeneity in cancer. Nature. 2022;611:810–7.

54. Dohlman AB, Klug J, Mesko M, Gao IH, Lipkin SM, Shen X, et al. A pan-cancer mycobiome analysis reveals fungal involvement in gastrointestinal and lung tumors. Cell. Elsevier; 2022;185:3807–3822.e12.

55. Fan AC, Thota RR, Serwas N, Vykunta VS, Marchuk K, Ruhland MK, et al. Submicrometre sampling of living cells by macrophages. Nature. Nature Publishing Group; 2026;654:495–503.

56. Roberts EW, Broz ML, Binnewies M, Headley MB, Nelson AE, Wolf DM, et al. Critical Role for CD103+/CD141+ Dendritic Cells Bearing CCR7 for Tumor Antigen Trafficking and Priming of T Cell Immunity in Melanoma. Cancer Cell. Elsevier; 2016;30:324–36.

57. Sanderson S, Shastri N. LacZ inducible, antigen/MHC-specific T cell hybrids. Int Immunol. 1994;6:369–76.

58. Gilles J-F, Dos Santos M, Boudier T, Bolte S, Heck N. DiAna, an ImageJ tool for object-based 3D co-localization and distance analysis. Methods. 2017;115:55–64.

