## Supplemental Figures for "Self-antigen disrupts cDC1 mediated antitumor responses"

#### Supplemental Figure 1.

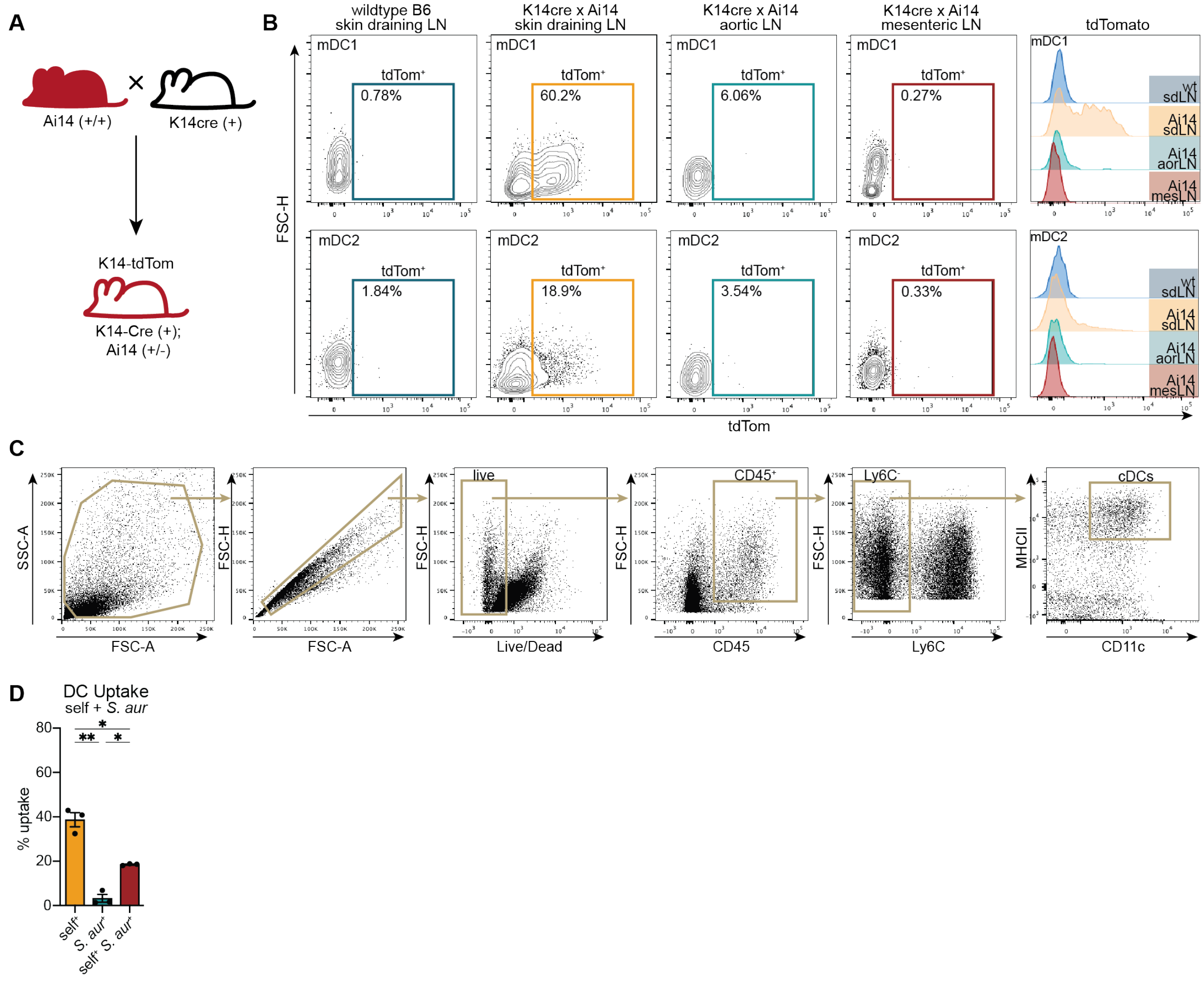


**Figure S1. Validation of mouse line and cDC uptake and antigen distribution.**
(A) Schematic of generation of K14-tdTom mouse line.
(B) Representative flow cytometry plots for tdTom+ migratory cDC1s (top) and cDC2s (bottom) from draining (skin LN) or non-draining (aortic, mesenteric LNs) lymph nodes. Data shown is representative of two biological replicates.
(C) Gating strategy for (Live/CD45+/Ly6C-/MHCII+CD11c+) cDCs from skin.
(D) Skin cDC uptake quantification of *S. aur*-mCh colonized K14-Cre; lsl-ZsG. Each point represents a single independent experiment, with pooled dorsal skin tissue from 6-12 mice per experiment. Data are shown as mean ± SEM. Statistical analysis was performed using RM one-way ANOVA, Geisser-Greenhouse correction, with Tukey's multiple comparison test. $*p < 0.05$, $**p < 0.01$, $***p < 0.001$, $****p < 0.0001$.

**Supplemental Figure 2.**

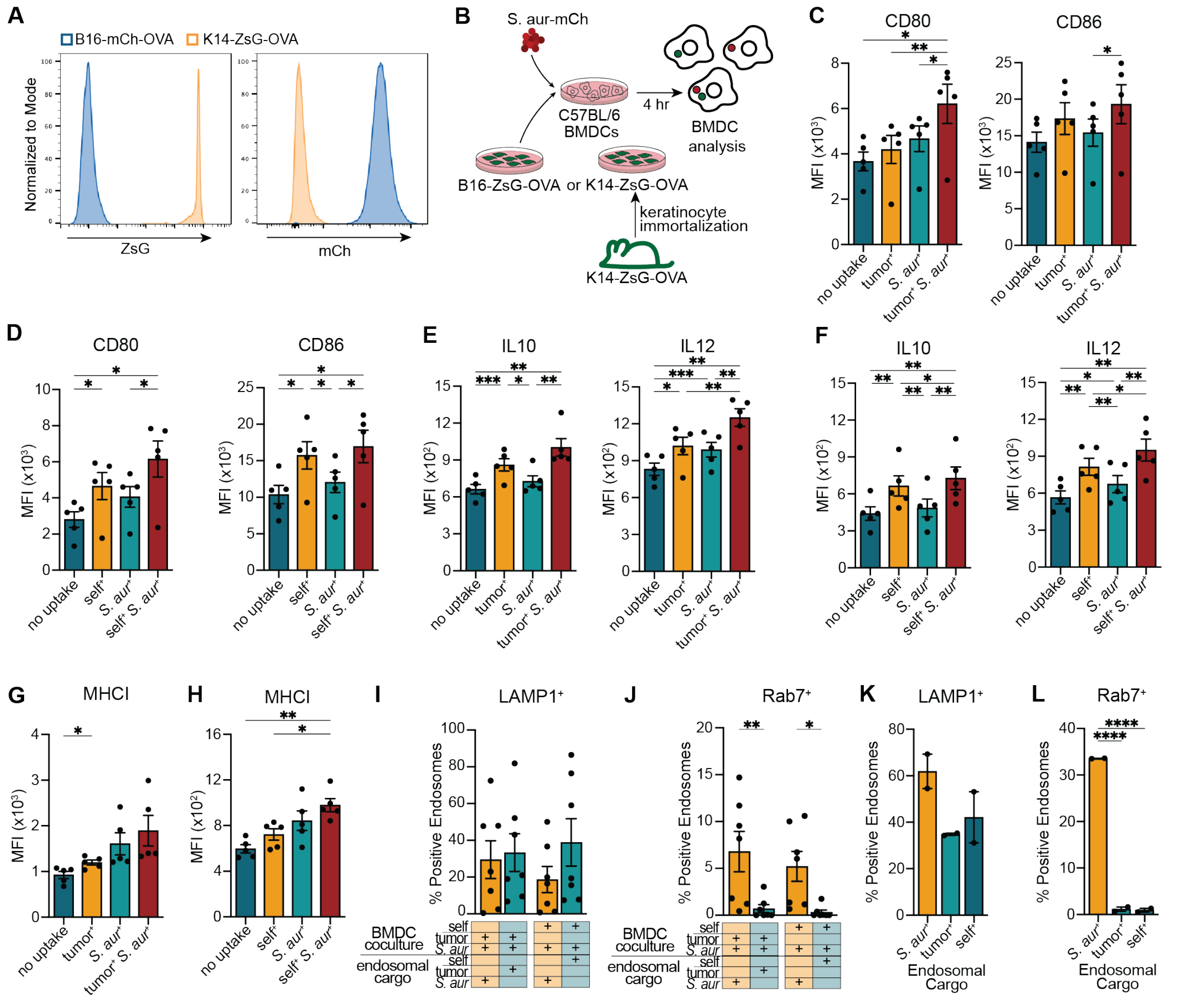

**Figure S2. Dual antigen uptake impact on BMDC maturation and endosomal processing.**
(A) Representative flow cytometry plot of B16-mChOVA and K14-ZsG-OVA cell lines.
(B) Schematic for cDC maturation from BMDC coculture with tumor (B16-ZsG-OVA) or self (K14-ZsG-OVA) and *S. aur*-mCh.
(C-D) MFI of CD80 (left) and CD86 (right) from surface staining of BMDCs cocultured with (C) tumor and *S. aur* or (D) self and *S. aur,* gated by uptake of indicated antigen.
(E-F) MFI of IL10 (left) and IL12 (right) from intracellular staining of BMDCs cocultured with (E) tumor and *S. aur* or (F) self and *S. aur,* gated by uptake of indicated antigen.
(G-H) MFI of surface MHCI from surface staining of BMDCs cocultured with (G) tumor and *S. aur* or (H) self and *S. aur,* gated by uptake of indicated antigen.
(I-J) Percent (I) LAMP1$^{+}$ (J) Rab7$^{+}$ of endosomes based on endosomal cargo from BMDCs pulsed with tumor and *S. aur* or self and *S. aur*.
(K-L) Percent (K) LAMP1$^{+}$ (L) Rab7$^{+}$ of endosomes from BMDCs pulsed with single antigen sources.

(C-H) Data are shown as mean ± SEM. Statistical analysis was performed using RM one-way ANOVA, Geisser-Greenhouse correction, with Tukey's multiple comparison test. Each point represents data from an individual mouse (n = 5, 3 independent experiments). (I-J) Data are shown as mean ± SEM. Statistical analysis was performed using RM two-way ANOVA with matched values. Post hoc comparisons were conducted using Fisher's LSD test. Each point represents data from an individual BMDC experiment (n = 4-7, 4-7 independent experiments). (K-L) Data are shown as mean ± SEM. Statistical analysis was performed using one-way ANOVA with Tukey's multiple comparison test. Each point represents data from an individual BMDC experiment (n = 2, 2 independent experiments). *$p < 0.05$, **$p < 0.01$, ***$p < 0.001$, ****$p < 0.0001$.

#### Supplemental Figure 3.

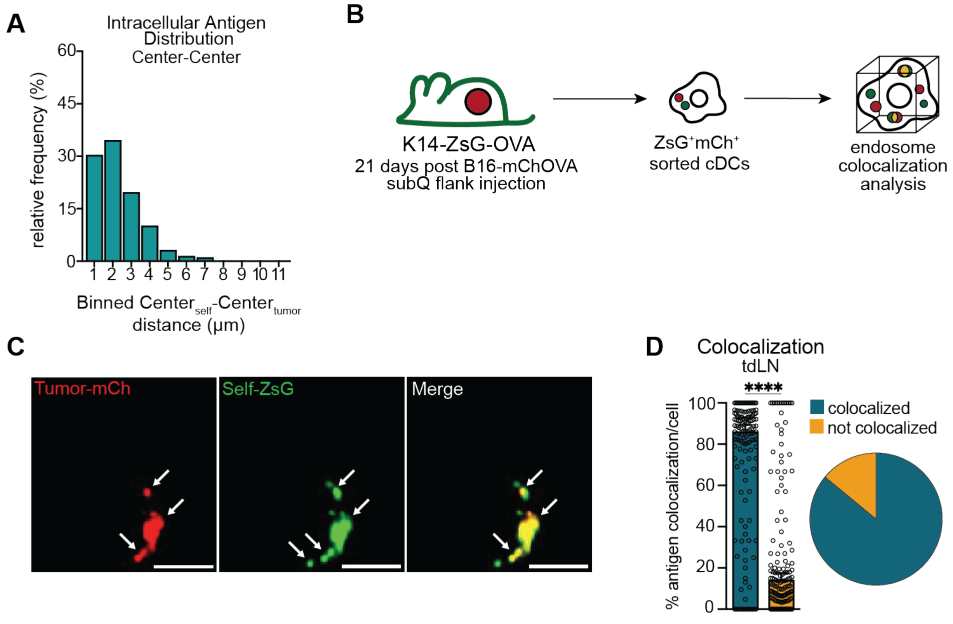


**Figure S3. cDC processing of self and tumor antigen is similar with tolerized ovalbumin.**
(A) Binned center-to-center distance analysis between intracellular antigens of each antigen source in tdLN. Data are shown as mean distance between CenterA- CenterB of antigens within an individual cell from (Fig. 3I).
(B)Schematic for colocalization analysis of tumor and self antigen by cDCs with OVA on both sources *in vivo*.
(C) Representative images of tumor$^{+}$self$^{+}$ cDCs from tdLN.
(D) Quantification of cDC colocalization of tumor-OVA and self-OVA antigen within the tdLN.

(D) Data are shown as median ± SEM. Statistical analysis was performed using a two-tailed unpaired Mann-Whitney U test. Each point represents the percentage of colocalized antigen within a single cell, 3 independent experiments with cells sorted from the pooled tissue of 8-12 mice. *p < 0.05, **p < 0.01, ***p < 0.001, ****p < 0.0001.

**Supplemental Figure 4.**

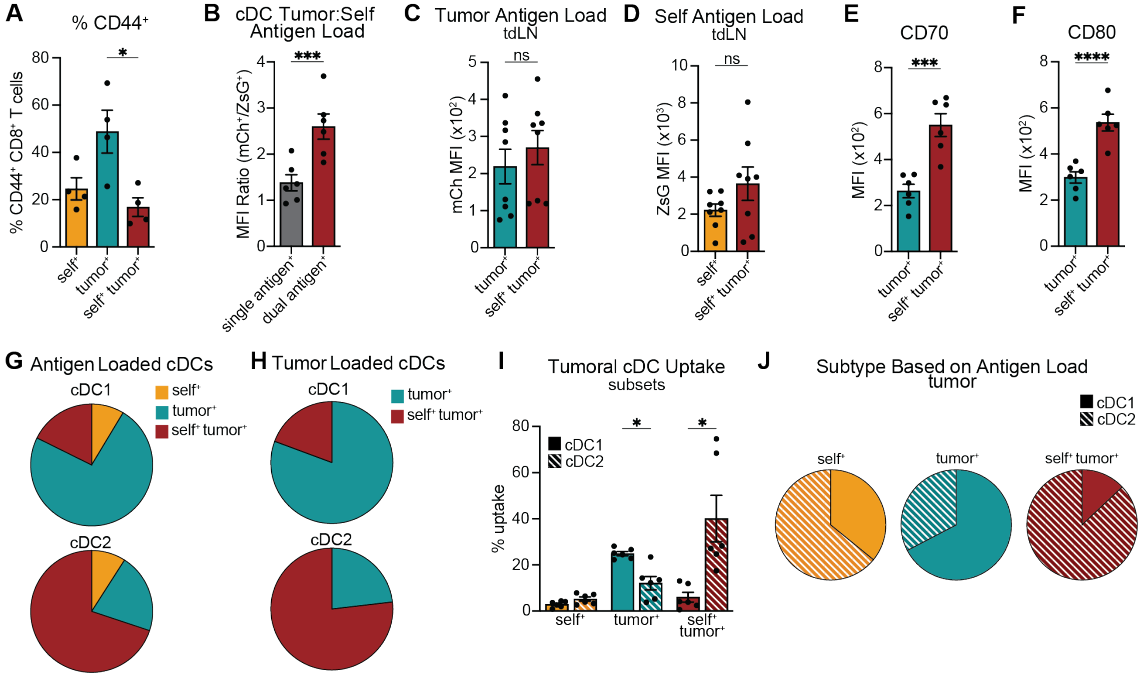


#### Figure S4. cDC antigen load and priming of tumor-specific CD8⁺ T cells upon tumor and self uptake.

(A) Quantification of percent CD44⁺ OT-Is cocultured with cDCs sorted based on uptake.
(B) Quantification of uptake by MFI ratio of mCh/ZsG of mCh⁺ cDCs:ZsG⁺ cDCs compared to MFI ratio of mCh/ZsG of ZsG⁺mCh⁺ cDCs:ZsG⁺mCh⁺ cDCs from tumor.
(C-D) Quantification of uptake by MFI of (C) tumor antigen or (D) self antigen by tdLN cDCs.
(E-F) MFI of (E) CD70 and (F) CD80 from tumor⁺, or tumor⁺self⁺ intratumoral DCs.
(G-H) ratio of uptake by (G) antigen⁺ or (H) tumor⁺ migratory cDC1s (mDC1s) and migratory cDC2s (mDC2s) from the tumor.
(I-J) (I) Quantified uptake and (J) Ratio of cDC subtype (mDC1 or mDC2) based on gating of antigen uptake by cDCs in the tumor.

(A) Data are shown as mean ± SEM. Statistical analysis was performed using one-way ANOVA, with Tukey's multiple comparison test. n = 4, 4 independent experiments.
(B-F) Data are shown as mean ± SEM. Statistical analysis was performed using two-tailed paired t-test. Each point represents an individual mouse. (B, E-F) n = 6 (C-D) n=8, 2 independent experiments.
(G-J) Data are shown as mean ± SEM. n = 6, 2 independent experiments. (I) Statistical analysis was performed using two-way RM ANOVA with matched values (stacked and across rows) and Geisser-Greenhouse correction. Post hoc comparisons were conducted using Šídák's multiple comparisons test. Each point represents an individual mouse. *$p < 0.05$, **$p < 0.01$, ***$p < 0.001$, ****$p < 0.0001$.

### Supplemental Figure 5.

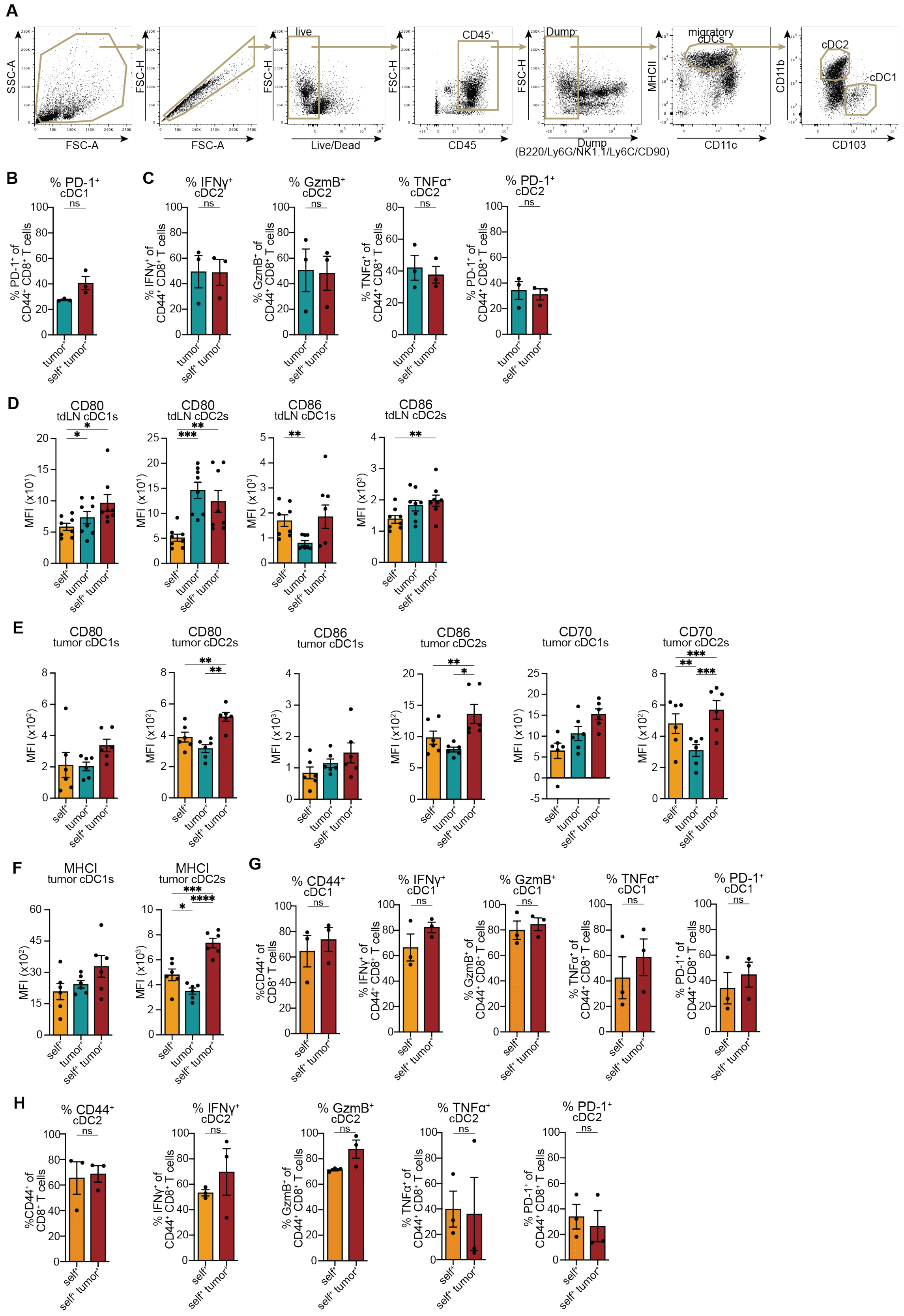

**Figure S5. Tumor$^{+}$self$^{+}$ cDC subset maturation and function of anti-tumor or anti-self primed CD8+ T cells from tumor$^{+}$self$^{+}$ cDC subsets.**

(A) Gating strategy for (Live/CD45$^{+}$/B220$^{-}$Ly6G$^{-}$NK1.1$^{-}$Ly6C$^{-}$CD90$^{-}$/MHCII$^{+}$CD11c$^{+}$) migratory cDC1 and cDCs from tdLNs.

(B-C, G-H) Percent (G-H) CD44$^{+}$, (C, G-H) IFNg$^{+}$, GzmB$^{+}$, TNFa$^{+}$ (B-C, G-H) PD-1$^{+}$ of (B-C) tumor-specific or (G-H) self-specific OT-Is following coculture with (B, G) mDC1s or (C, H) mDC2s that are (B-C) tumor$^{+}$ or tumor$^{+}$self$^{+}$ or (G-H) self$^{+}$ or tumor$^{+}$self$^{+}$.

(D-F) MFI of (D-E) CD80, CD86, (E) CD70, (F) MHCI of mDC1s (left) and mDC2s (right) based on antigen uptake in (D) tdLN or (E-F) tumor.

(B-C, G-H) Data are shown as mean ± SEM. Statistical analysis was performed using two-tailed paired t-test. Each point represents the mean data from an individual experiment set up with 14-18 mice pooled per experiment. n = 3 independent experiments.

(D-F) Data are shown as mean ± SEM. (D) n = 8, (F) n = 6, 2 independent experiments. Each point represents an individual mouse. Statistical analysis was performed using RM one-way ANOVA, Geisser-Greenhouse correction, with Tukey's multiple comparison test. $*p < 0.05$, $**p < 0.01$, $***p < 0.001$, $****p < 0.0001$.

**Supplemental Figure 6.**

A

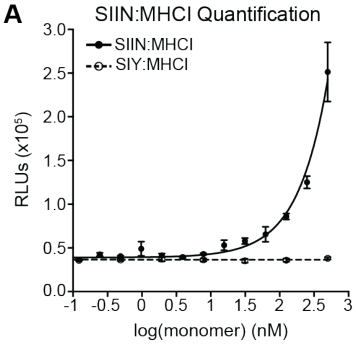


**Figure S6. Standard curve of B3Z stimulation assay.**

(A) Standard curve of either SIIN:MHCI or SIY:MHCI monomers titrated against B3Z cells in luminescent assay with RLUs. Mean ± SEM. n = 3 per titrated data point. Nonlinear regression of exponential growth equation with RLU constraints for Y0 = mean RLUs without monomer.
